# Human liver microbiota modeling strategy at the early onset of fibrosis

**DOI:** 10.1101/2020.12.10.419051

**Authors:** Camille Champion, Radu M. Neagoe, Maria Effernberger, Daniela T. Sala, Florence Servant, Jeffrey E. Christensen, Maria Arnoriaga-Rodriguez, Jacques Amar, Benjamin Lelouvier, Fabrice Gamboa, Herbert Tilg, Massimo Federici, Jose-Manuel Fernández-Real, Jean Michel Loubes, Rémy Burcelin

**Affiliations:** Institut National de la Santé et de la Recherche Médicale (INSERM), Toulouse, France; Université Paul Sabatier (UPS), Unité Mixte de Recherche (UMR) 1048, Institut des Maladies Métaboliques et Cardiovasculaires (I2MC), Team 2 : ‘Intestinal Risk Factors, Diabetes, Dyslipidemia’ F-31432 Toulouse Cedex 4, France; Institut de Mathématiques de Toulouse, Université Paul Sabatier, Toulouse, France; University of Medicine Pharmacy, Science and Technology “George Emil Palade” Tîrgu Mures, Second Department of Surgery, Emergency Mureş County Hospital, Romania; Department of Internal Medicine I, Gastroenterology, Hepatology, Endocrinology & Metabolism, Medical University of Innsbruck, Austria; VAIOMER, 516 Rue Pierre et Marie Curie, 31670 Labège, France; Department of Diabetes, Endocrinology and Nutrition, University Hospital of Girona ‘Dr Josep Trueta’; Institut d’Investigacio Biomedica de Girona IdibGi; and CIBER Fisiopatologia de la Obesidad y Nutricion. Girona, Spain; Therapeutics department, Rangueil Hospital, Toulouse, France; Department of Systems Medicine, University of Rome “Tor Vergata”, Via Montpellier 1, 00133 Rome, Italy

**Author notes:** Corresponding authors and Lead Contact: Pr Remy Burcelin; Pr Radu Mircea Neagoe, and Pr jean Michel Loubes. contributed equally to this work (Camille Champion, Radu M. Neagoe).

**Keywords:** biomathematics, liver diseases, metabolic disease, microbiota, tissue microbiota

## Abstract

To understand the pathophysiological impact of liver microbiota on the early stages of fibrosis we identified the corresponding microbiota sequences and overcome the impact of different group size and patient origins with adapted statistical approaches. Liver samples with low liver fibrosis scores (F0, F1, F2) were collected from Romania(n=36), Austria(n=10), Italy(n=19), and Spain(n=17). The 16SrDNA gene was sequenced. We considered the frequency, sparsity, unbalanced sample size between cohorts to identify taxonomic profiles and statistical differences. Multivariate analyses, including adapted spectral clustering with L1-penalty fair-discriminant strategies, and predicted metagenomics were used to identify that 50 % of liver taxa were Enterobacteriaceae and Pseudomonadaceae. The Caulobacteraceae, Flavobacteriaceae and Propionibacteriaceae discriminated between F0 and F1. The preQ0 biosynthesis and pathways involving glucoryranose and glycogen degradation were negatively associated with liver fibrosis F1-F2 vs F0. Altogether, our results suggest a role of bacterial translocation to the liver in the progression of fibrosis. This statistical approach can identify microbial signatures and overcome issues regarding sample size differences, the impact of environment, and sets of analyses.

## Introduction

Non-alcoholic fatty liver disease (NAFLD) is a common consequence of obesity and type 2 diabetes [1,2]. In NAFLD, the origin of inflammation and hepatocyte injury is related to dietary lipids, bile acids, adipokines and cytokines, to cite a few. Furthermore, gut microbiota seems to be one of the key players of NAFLD development [3,4]. Markers and receptors of microbiota-related injury features have been described in this disorder such as TLRs, NODS, and NLRP3 [5-8] as well as the activation of the innate and adaptive immune systems [9]. In early sets of experiments, we initially showed that hepatic steatosis in the obese diabetic mouse was due to an increased circulating concentration of lipopolysaccharides (LPS) i.e. metabolic endotoxemia [10]. Lipoproteins transport LPS [11] to tissues, triggering the CD14/TRL4 pathway that increases liver inflammation and fat deposition [10]. Gut bacteria were also reported to translocate through the intestinal tract to tissues [12] such as the adipose depots and the liver, establishing a tissue microbiota as observed in rodents [13-15] and humans [16-18] which could trigger liver inflammation and the onset of fibrosis [13]. This mechanism activates immune cells, including Kupffer cells, to release various pro-inflammatory cytokines and chemokines [19] damaging the surrounding tissues initiating fibrosis. This hypothesis is now largely supported by recent major advances in NAFLD research, which show gut and blood microbiota dysbiosis of patients with advanced stages of NAFLD [20-22]. Hence, the identification of specific groups of translocated bacteria from dysbiotic gut microbiota could aid in the design of novel therapeutic strategies. To address this issue, we have sequenced and identified the bacterial 16S rDNA from liver tissue of a cohort of 36 Romanian, 17 Spanish, 19 Italians and 10 Austrian patients with different stages of liver fibrosis, notably at their early stages. We could design hypotheses regarding the putative causal role of liver microbiota in the development of liver fibrosis. We used this database to evaluate the efficacy of Principal Coordinate Analysis (PCoA) to visualize the different liver fibrosis group scores using Wilcoxon-Mann-Whitney statistical tests [23]. Eventually, since the overall database of patients issued from different separated cohorts we anticipated some degree of heterogeneity of the overall cohort therefore, we adapted and developed a specific statistical approach i.e. L1 spectral clustering with fairness. This approach establishes inter-relations between liver microbiota and low scores of liver fibrosis that allowed the identification of the translocated bacteria putatively causal to the disease and independent from the group size, the patient origins and sets of sequencing. Overall, we drew a European microbial profile of patients at early stages of liver fibrosis.

## Materials and Methods

### Subjects and Ethics

A multicentric observational study was conducted in the Second Department of Surgery, Emergency Mureş County Hospital of Romania, the Department of Systems Medicine of the Tor Vergata University of Rome, the Institut d’Investigacio Biomedica de Girona IdibGi, the Endocrinology and Nutrition Department of Dr. Josep Trueta University Hospital, and the University Hospital of Innsbruck. All research procedures performed in this study were in strict accordance with a pre-defined protocol and adhered to the Good Clinical Practice guidelines and the Declaration of Helsinki. The study was approved by the Coordinating Ethics Committee of the Emergency Mureş County Hospital, Romania (registration 4065/2014), the Institutional review board & Ethics Committee and the Committee for Clinical Research (CEIC) of Dr. Josep Trueta University Hospital, Girona, Spain; the Policlinico Tor Vergata Ethics Committee, Rome, Italy as part of the FLORINASH Study the Institutional Ethics Commission at the medical University of Innsbruck (amendment to AN20170016 369/4.21). All participants provided informed consent prior to participation. The patients who gave their consent to perform a liver biopsy during the procedure were eligible. Exclusion criteria were serious liver diseases (eg hemochromatosis, alcoholic fatty liver disease, Hepatits B and Hepatitis C infection, chronic diseases, inflammatory systemic diseases, acute or chronic infections in the previous month, use of antibiotic, antifungal, antiviral drugs, proton-pump inhibitors, anti-obesity drugs, laxatives, excessive use of vitamin D supplementation, fiber supplements or probiotics or participation in a weight loss program or weight change of 3 kg during the previous 6 weeks, pregnancy or breastfeeding, or major psychiatric antecedents; neurological diseases, history of trauma or injured brain, language disorders, and excessive alcohol intake (≥ 40 g/day in women or 80g OH/day in men) or intravenous drug abuse, and previous bariatric surgery.

The cohort consists of 82 Caucasian patients where 34 were diagnosed with fibrosis stage 0 (F0); 37 stage 1 (F1) and 11 stage 2 (F2), as diagnosed from histological analyses of liver biopsies (**Table 1**). The patients suffered from morbid obesity with a mean BMI 42.6 (±7.3). The mean waist circumference was 121.49 (±18.73) in male and 123.23 (±18.26) in female participants.

**Table 1.**
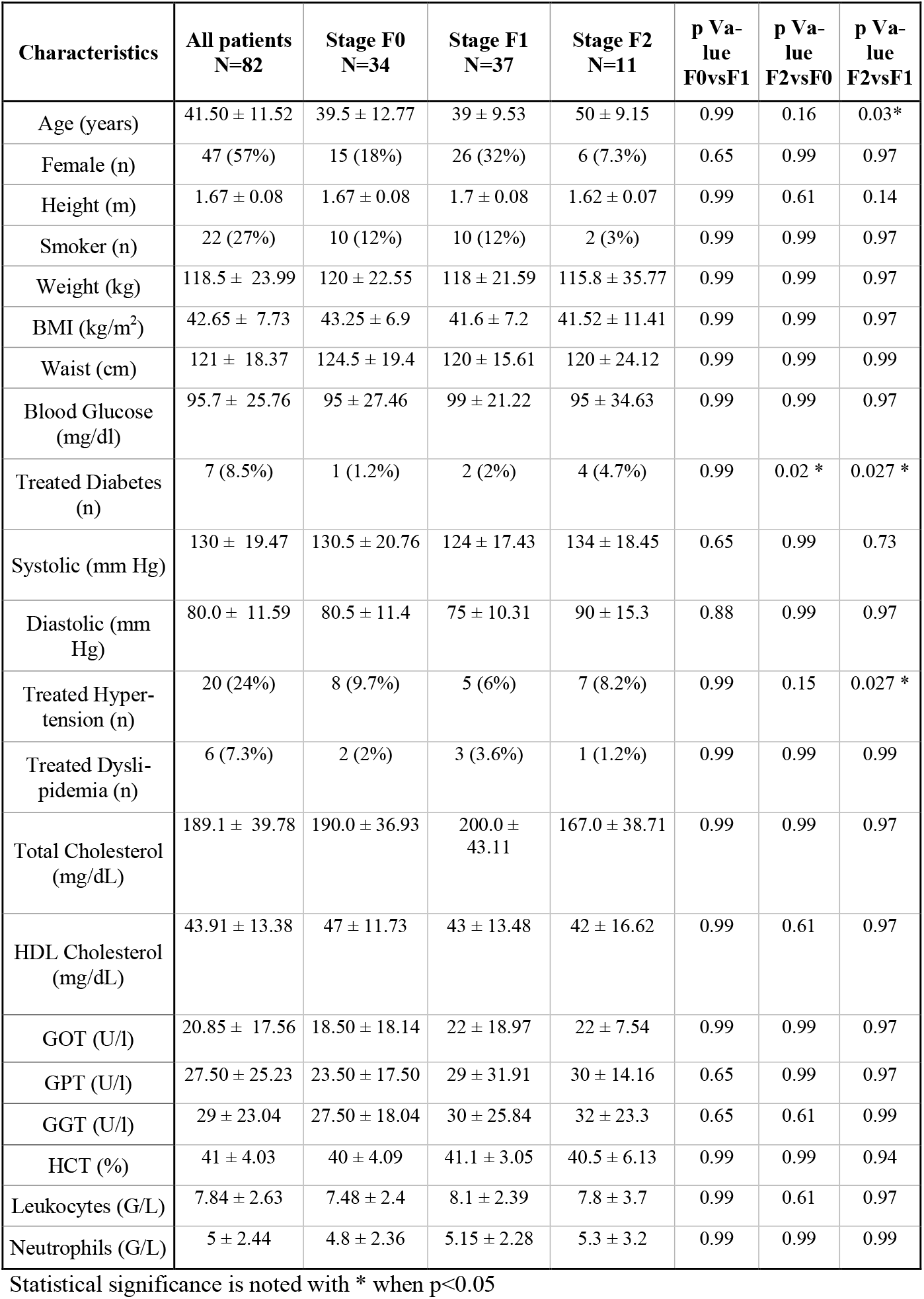
Baseline characteristics of patients with biopsy-proven fibrosis.

| Characteristics | All patients<br>N=82 | Stage F0<br>N=34 | Stage F1<br>N=37 | Stage F2<br>N=11 | p Value<br>F0vsF1 | p Value<br>F2vsF0 | p Value<br>F2vsF1 |
| --- | --- | --- | --- | --- | --- | --- | --- |
| Age (years) | 41.50 ± 11.52 | 39.5 ± 12.77 | 39 ± 9.53 | 50 ± 9.15 | 0.99 | 0.16 | 0.03* |
| Female (n) | 47 (57%) | 15 (18%) | 26 (32%) | 6 (7.3%) | 0.65 | 0.99 | 0.97 |
| Height (m) | 1.67 ± 0.08 | 1.67 ± 0.08 | 1.7 ± 0.08 | 1.62 ± 0.07 | 0.99 | 0.61 | 0.14 |
| Smoker (n) | 22 (27%) | 10 (12%) | 10 (12%) | 2 (3%) | 0.99 | 0.99 | 0.97 |
| Weight (kg) | 118.5 ± 23.99 | 120 ± 22.55 | 118 ± 21.59 | 115.8 ± 35.77 | 0.99 | 0.99 | 0.97 |
| BMI (kg/m<sup>2</sup>) | 42.65 ± 7.73 | 43.25 ± 6.9 | 41.6 ± 7.2 | 41.52 ± 11.41 | 0.99 | 0.99 | 0.97 |
| Waist (cm) | 121 ± 18.37 | 124.5 ± 19.4 | 120 ± 15.61 | 120 ± 24.12 | 0.99 | 0.99 | 0.99 |
| Blood Glucose<br>(mg/dl) | 95.7 ± 25.76 | 95 ± 27.46 | 99 ± 21.22 | 95 ± 34.63 | 0.99 | 0.99 | 0.97 |
| Treated Diabetes<br>(n) | 7 (8.5%) | 1 (1.2%) | 2 (2%) | 4 (4.7%) | 0.99 | 0.02 * | 0.027 * |
| Systolic (mm Hg) | 130 ± 19.47 | 130.5 ± 20.76 | 124 ± 17.43 | 134 ± 18.45 | 0.65 | 0.99 | 0.73 |
| Diastolic (mm<br>Hg) | 80.0 ± 11.59 | 80.5 ± 11.4 | 75 ± 10.31 | 90 ± 15.3 | 0.88 | 0.99 | 0.97 |
| Treated Hyper-<br>tension (n) | 20 (24%) | 8 (9.7%) | 5 (6%) | 7 (8.2%) | 0.99 | 0.15 | 0.027 * |
| Treated Dysli-<br>pidemia (n) | 6 (7.3%) | 2 (2%) | 3 (3.6%) | 1 (1.2%) | 0.99 | 0.99 | 0.99 |
| Total Cholesterol<br>(mg/dL) | 189.1 ± 39.78 | 190.0 ± 36.93 | 200.0 ± 43.11 | 167.0 ± 38.71 | 0.99 | 0.99 | 0.97 |
| HDL Cholesterol<br>(mg/dL) | 43.91 ± 13.38 | 47 ± 11.73 | 43 ± 13.48 | 42 ± 16.62 | 0.99 | 0.61 | 0.97 |
| GOT (U/l) | 20.85 ± 17.56 | 18.50 ± 18.14 | 22 ± 18.97 | 22 ± 7.54 | 0.99 | 0.99 | 0.97 |
| GPT (U/l) | 27.50 ± 25.23 | 23.50 ± 17.50 | 29 ± 31.91 | 30 ± 14.16 | 0.65 | 0.99 | 0.97 |
| GGT (U/l) | 29 ± 23.04 | 27.50 ± 18.04 | 30 ± 25.84 | 32 ± 23.3 | 0.65 | 0.61 | 0.99 |
| HCT (%) | 41 ± 4.03 | 40 ± 4.09 | 41.1 ± 3.05 | 40.5 ± 6.13 | 0.99 | 0.99 | 0.94 |
| Leukocytes (G/L) | 7.84 ± 2.63 | 7.48 ± 2.4 | 8.1 ± 2.39 | 7.8 ± 3.7 | 0.99 | 0.61 | 0.97 |
| Neutrophils (G/L) | 5 ± 2.44 | 4.8 ± 2.36 | 5.15 ± 2.28 | 5.3 ± 3.2 | 0.99 | 0.99 | 0.99 |

Statistical significance is noted with \* when p<0.05

### Liver biopsies and liver fibrosis diagnosis

Liver biopsies were performed during laparoscopic surgical bariatric procedures or via ultrasound guided liver biopsy. No energy devices were used for collecting the samples since hemostasis was done afterwards when the samples were extracted from the abdomen. Ultrasound (US) guided percutaneous liver biopsy (UPLB) was performed in 10 patients. In all patients, antiplatelet drugs and oral anticoagulation therapy was paused 1 week before UPLB was performed. One experienced physician (> 3000 US-exams and >100 UPLB) performed the US-examinations with the Philips EPIQ 5^®^ (Philips Corporation, Amsterdam, The Netherlands). UPLB was performed using an 18 G Temno II semi-automatic tru-cut biopsy needle (Cardinal Health, Dublin, Ohio, USA). After UPLB, all patients were monitored for any signs of pain or clinically suspected bleeding by nursing staff over a 6-h period. If no serious complications were evident, all patients would be discharged after the mandatory 6-h observation, a stable blood count and a normal ultrasound examination. All patients were follow-up in 2 weeks to review the results of the histology. All the samples were stored in a sterile container and kept at −80°C until assayed. Furthermore, NAFLD was confirmed histologically by an independent pathologist.

### Clinical assessments

Anthropometric measurement of each subject was performed by trained nurses in the morning after fasting for at least 8 h. Body height was recorded to the nearest 0.5 cm and body weight to the nearest 0.1 kg. BMI was defined as body weight (kilograms) divided by the square of body height (meters). Waist circumference was measured in the horizontal plane midway between lowest rib and the iliac crest to the nearest 0.1 cm at the end of a normal expiration repeatedly in men and women by 3 trained nurses on 3 consecutive days. Blood pressure was recorded to the nearest 2 mmHg by a mercury sphygmomanometer with the arm supported at heart level after sitting quietly for 10 min. Fasting plasma glucose was measured after fasting for at least 8 h. A standard oral 75-g glucose tolerance test was performed to measure 2-h postprandial plasma glucose. Hypertension was defined in accordance to the Guidelines of the European Heart Association or if the subject was taking medication for hypertension. Diabetes was diagnosed when fasting plasma glucose was ≥126 mg/dL (7 mmol/L), 2-h postprandial plasma glucose ≥200 mg/dL (11.1 mmol/L), and HbA_1c_ ≥6.5% or if the subject was taking medication for diabetes

### Biochemical and molecular analyses

#### Plasma parameters

Biochemical analyses including total fasted plasma glucose, cholesterol, high-density lipoprotein (HDL) cholesterol, plasma liver enzymes i.e. aspartate aminotransferase (AST/GOT), alanine aminotransferase (ALT/GPT), gamma-glutamyl transferase (GGT), hematocrit and leukocytes were determined by Cobas 8000, (Roche, Basel, Switzerland) according to the manufacturer’s specification. Elevated liver enzymes were defined as aspartate aminotransferase and alanine aminotransferase. HbA1c was measured by high-performance liquid chromatography (Bio-Rad, Muenchen, Germany) and a Jokoh HS-10 autoanalyzer.

#### 16S rDNA sequencing and bioinformatic analysis

The V3-V4 hypervariable regions of the 16S_rDNA were amplified by two steps PCR using v1 primers (Vaiomer) and sequenced using MiSeq Reagent Kit v3 (2×300 bp Paired-End Reads, Illumina, San Diego, CA, USA) as previously described [24]. The MiSeq sequences were then analyzed using the bioinformatics pipeline established by Vaiomer using FROGS v1.4.0 [25]. Briefly, after demultiplexing of the bar-coded Illumina paired reads; single read sequences are cleaned and paired for each sample independently into longer fragments. Operational taxonomic units (OTU) are produced with via single-linkage clustering and taxonomic assignment is performed in order to determine community profiles (generated by Blast+ v2.2.30+ against the Silva v128 Parc databank restricted to the bacterial kingdom).

#### Linear Discriminant Analysis (LDA) Effective Size (LEfSe)

The bacterial profiles were further compared between the three groups using LEfSe pairwise analysis with an alpha cut-off of 0.05 and an effect size cut-off of 2.0. The bacterial diversity analyses (alpha and beta diversity, MDS ordinations and taxonomic composition barplots) were generated using the Phyloseq (v1.14.0), vegan (v2.4.0) and ape (v3.5) packages under R environment v3.3.1. LEfSe analysis was performed on the OTU table using the online Galaxy interface to identify bacterial taxa that were differentially abundant in the three liver fibrosis groups [27]. Respective cladograms were generated with genus at the lowest level. Quantitative plots of differential features were generated from genus level percent relative abundance data showing means with standard deviation using GraphPad Prism 6 software. Using the LEfSe algorithm, bacterial taxa that were differentially abundant in analysis of liver fibrosis groups were first identified and tested using the Kruskal Wallis test.

#### Beta diversity analysis

The bacterial diversity (alpha and beta diversity) was analyzed and represented using the phyloseq (v1.14.0), vegan (v2.4.0), ape (v3.5), and ggplot (3.3.0) packages under R environment v3.5.1 with Chao, Inverse Simpson, Simpson and Shannon as indexes. The alpha diversity statistical significance was determined by Wilcoxon rank-test. The beta diversity was calculated for every pair of variables to generate a matrix of distance using Bray-Curtis, Jaccard, Unifrac, and weighted Unifrac indexes. From distance matrices, Multiple Dimension Scale (MDS) and hierarchical clustering were conducted for graphical representation of beta diversity. PERMDISP2 procedure was used for the analysis of multivariate homogeneity of group dispersions. The Kruskall-Wallis test was performed to compare abundance across the three groups.

#### Multivariate analyses

To visualize the distribution of patients according to their clinical parameters, we performed a Principal Component Analysis (PCA) using FactoMineR and factoextra R packages. For the study of 16SrDNA diversity, we first filtered the less abundant OTUs to reduce the noise within the matrix before running the PCA. We eliminated those with abundance <0.01. We then normalized the OTU table by using the Cumulative Sum Scaling normalization followed by a log transformation, using mixOmics package (https://pubmed.ncbi.nlm.nih.gov/29099853/). To explore the metagenomic data and identify the largest sources of variation, another Principal Component Analysis was conducted. Also based on the projection of the dataset into a space of lower dimension and originally designed for regression we performed a Partial Least Square Discriminant Analysis (PLS-DA) and its sparse version (sPLS-DA) on the normalized OTU table count to predict and select the most discriminative features in the data that help to classify the samples according to the fibrosis variable (package mixOmics).

Since we observed the influence of the metagenomic data on the outcome, we used alternative method of classification such as random forest (package randomForest). The random forest is built from a multitude of different decision trees and classifiers at training time thereby predicting and storing the predicted target outcome.

Cluster graphical analyses. The abundance matrix of OTUs can be modeled by a graph using PLNmodels package under R where nodes represent OTUs and edges interactions between each pair of nodes. We developed an analysis in clusters i.e. the L1-spectral clustering, implemented in R, a robust variant of the well-known spectral clustering that aims to detect the natural structures of a graph by taking advantage of its spectral properties. The adjacency matrix modeling the variable associations of the graph is used as an input of the l1-spectralclustering algorithm. In front of the influence of the origin of the cohort on the graphical classification through clusters we applied “fair” technics with k-median clustering objectives. We identified k centers and assign each input point to one of the centers so that the average distance of points to their cluster center is minimized. In the fair-variant, the points are colored while the goal is to minimize the same average distance objective ensuring all clusters to have an approximately equal number of points of each color. This technique called “fairtree” and developed in python takes as input the desired number of clusters, the desired cluster balance and the normalized table count.

#### Functional metagenomic prediction

Metagenome inference and predicted functional analysis were initiated by analysis of the OTU clustered 16S sequence count table data and the OTU representative sequences using the PICRUSt2 tool [26] (**https://pubmed.ncbi.nlm.nih.gov/32483366/)** version 2.3.0b for each sample. The metagenome prediction process included four main steps: 1) The input OTU representative sequences were aligned against the PICRUSt2 reference alignment, 2) From this alignment, the input OTU were placed into the PICRUSt2 reference phylogenetic tree, 3) The metagenome functions were inferred by the hidden state prediction method using this phylogenetic tree. During this inference process, the abundance values of each OTU were normalized to their respective predicted 16S_rDNA copy numbers and then multiplied by the respective gene counts of the target bacteria, 4) The predicted functions were mapped to the MetaCyc database to determine the minimum set of pathways present in the samples. The resulting core output was a list of enzyme functions (Enzyme Commission numbers) with predicted count data for each sample from step 3 as well as a list of MetaCyc pathways with predicted count data for each sample from step 4.

### Data Availability Section

- MiSeq 16S_rDNA sequences were deposited under the primary accession number PRJEB41831 and a secondary number ERP125667 on December 9^th^ 2020 with a release date on the 31^st^ of December 2021.

## Results

### Graphical classification of the clinical variables by principal component analyses

We aimed at identifying liver 16SrDNA profiles associated with the early onset of fibrosis. We aggregated together a library of liver biopsies from patients from four cohorts of different European countries. We first visualized the distribution of the patients according to the cohorts by performing a principal component analysis using the anthropomorphic and clinical data where the projection of the different clinical variables is represented (**Fig1A,B**). The ellipses calculated for each cohort show some degree of differential distribution suggesting that specific environmental factors have influenced the clinical outcomes. Interestingly, the Romania cohort was unifying all cohorts and could be used as a reference. In addition, we could detect numerous outlier patients from each cohort.

**Fig. 1:**
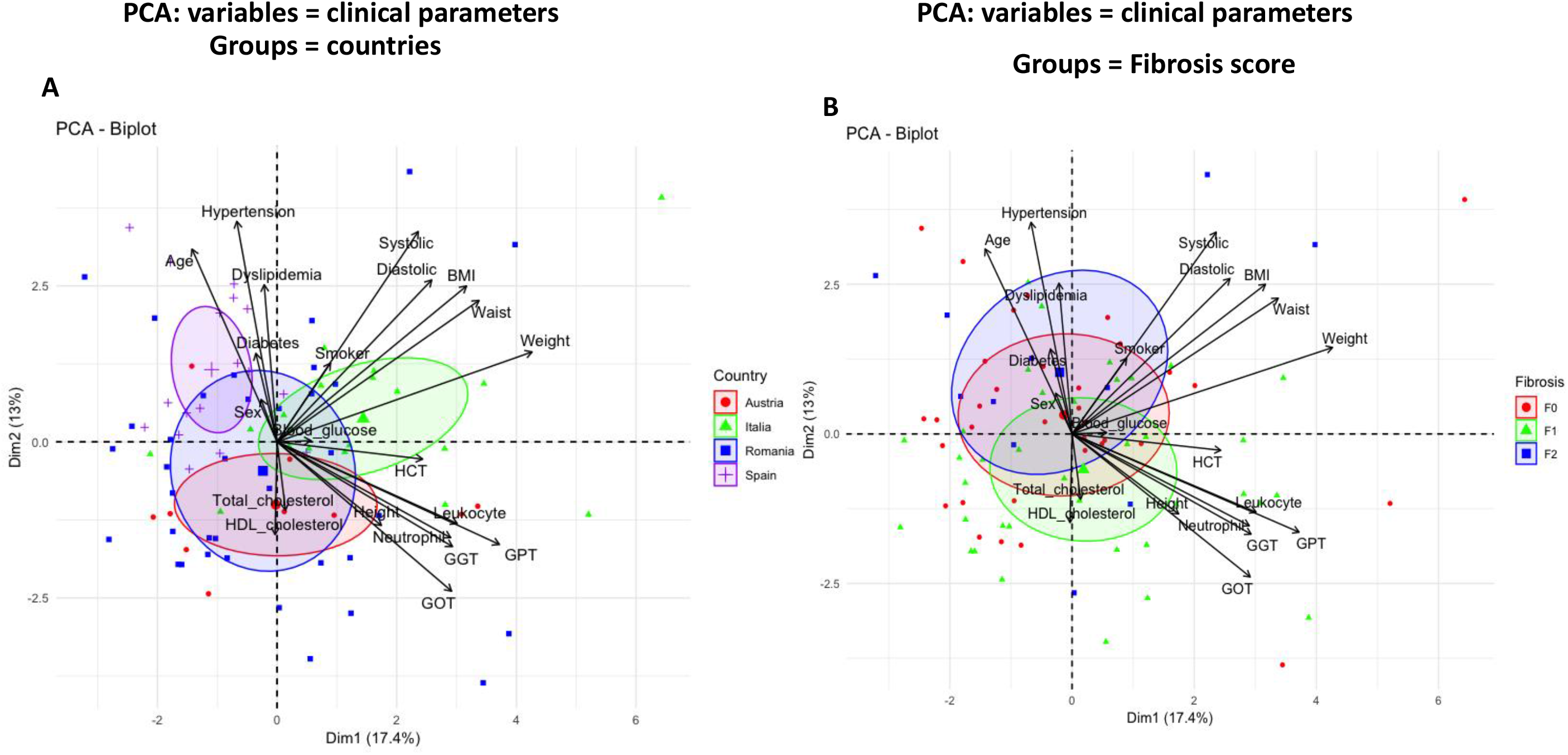
Visualization of clinical variables by principal component analysis according to countries and fibrosis scores. The clinical variables were used as entries for a principal component analysis (PCA). PCA-biplot from package Factoextra and FactomineR of individuals for the first two principal components are shown. They sum up 30.4% of the total variance of the dataset. Patients were grouped by **A**, countries (red dots=Austria, green triangle=Italy, blue square=Romania, purple cross=Spain) and by **B**, fibrosis scores (red dots=F0, green triangle=F1, blue square=F2). The vectors corresponding to the clinical variables are shown as arrows.

It is noteworthy that we voluntarily included all anthropomorphic and biochemical data, even if some were redundant and confounding, to remain within the frame of a non-a priory statistical approach. The age, diabetes and hypertension variables were the main drivers of the F2 classification while HDL cholesterol and liver enzymes were drivers for the F1 histological phenotype. These observations are supported by significant ANOVA tests (**Table 1**).

### Analyses of the liver bacterial 16SrDNA ecology

To identify whether the graphical differences between the three liver fibrosis scores are associated with a differential liver bacterial DNA signature, we then performed PCA on the OTUs as entries in the database. The analysis using countries as groups shows that the different cohorts poorly overlapped suggesting the existence of specific environmental factors specific of each country cohort (**Fig 2A**). Using the liver fibrosis scores as groups we could not clearly graphically discriminate the fibrosis scores since the distribution of the patients according to their OTU profiles were too scattered and seemed to be depending upon the largest Romanian cohort (**Fig 2B**). To analyze differently the putative signatures according to the cohorts or the liver fibrosis scores, we studied the frequencies of the phylum and family taxonomic levels. The barplot analysis shows first a large degree of heterogeneity between all individuals at the phylum level (**Fig 2C**) but still, we identified that the liver microbiota of the overall cohort was composed mostly of Proteobacteria, (>75%) (**Fig 2D**). Group comparisons showed that statistical differences were observed between the F0 and F1 groups for the Proteobacteria, Actinobacteria and Firmicutes phyla (**Supplementary Fig1 A**,**B**,**C**). At the family taxonomic level, the most prominent taxa were the Enterobacteriaceae and the Pseudomonadaceae which accounted for more than 50 % of the overall taxa (**Fig 2E**). Group comparisons showed that the Caulobacteraceae, Flavobacteriaceae and Propionibacteriaceae families were statistically different when comparing F0 and F1 (**Supplementary Fig1 D,E,F**).

**Fig. 2:**
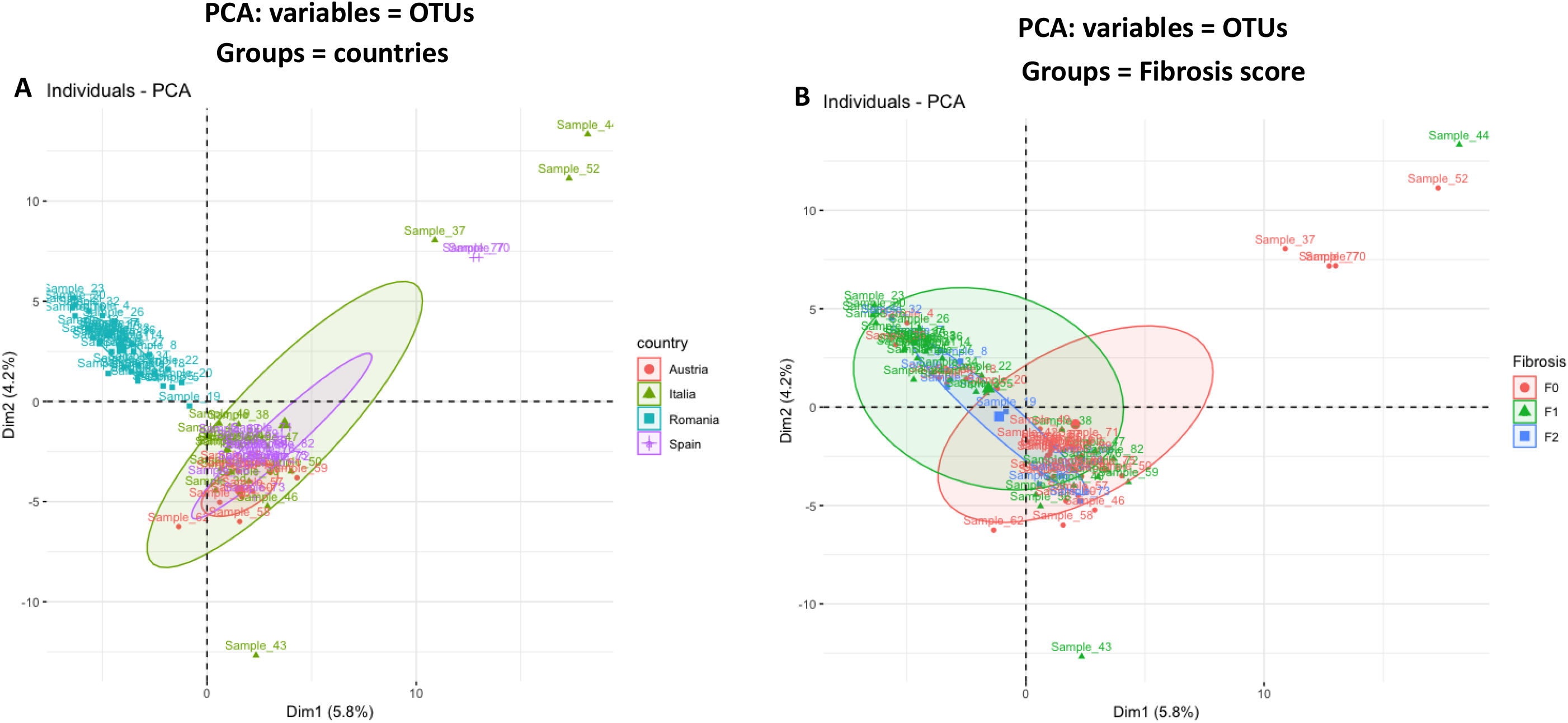
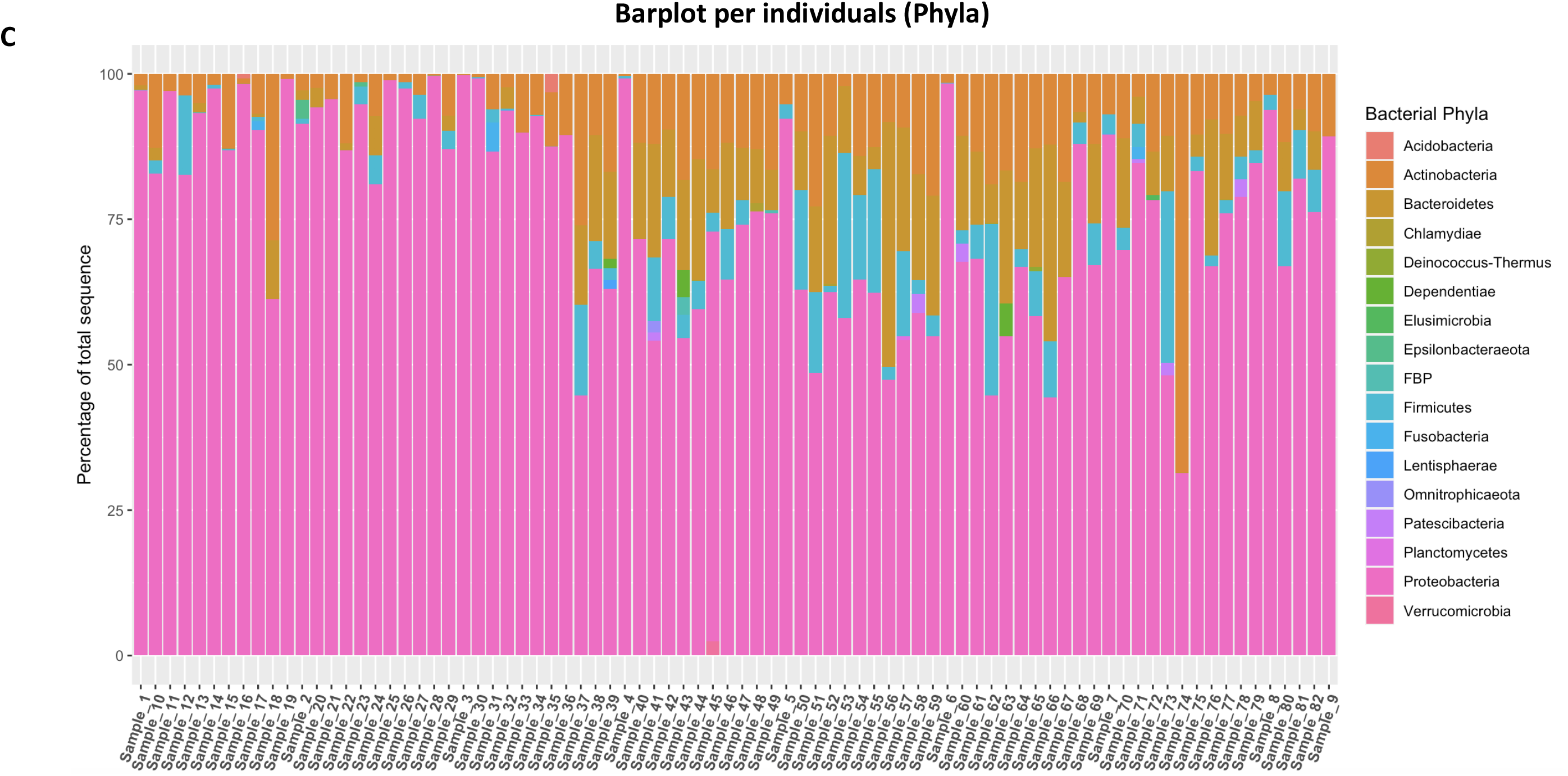
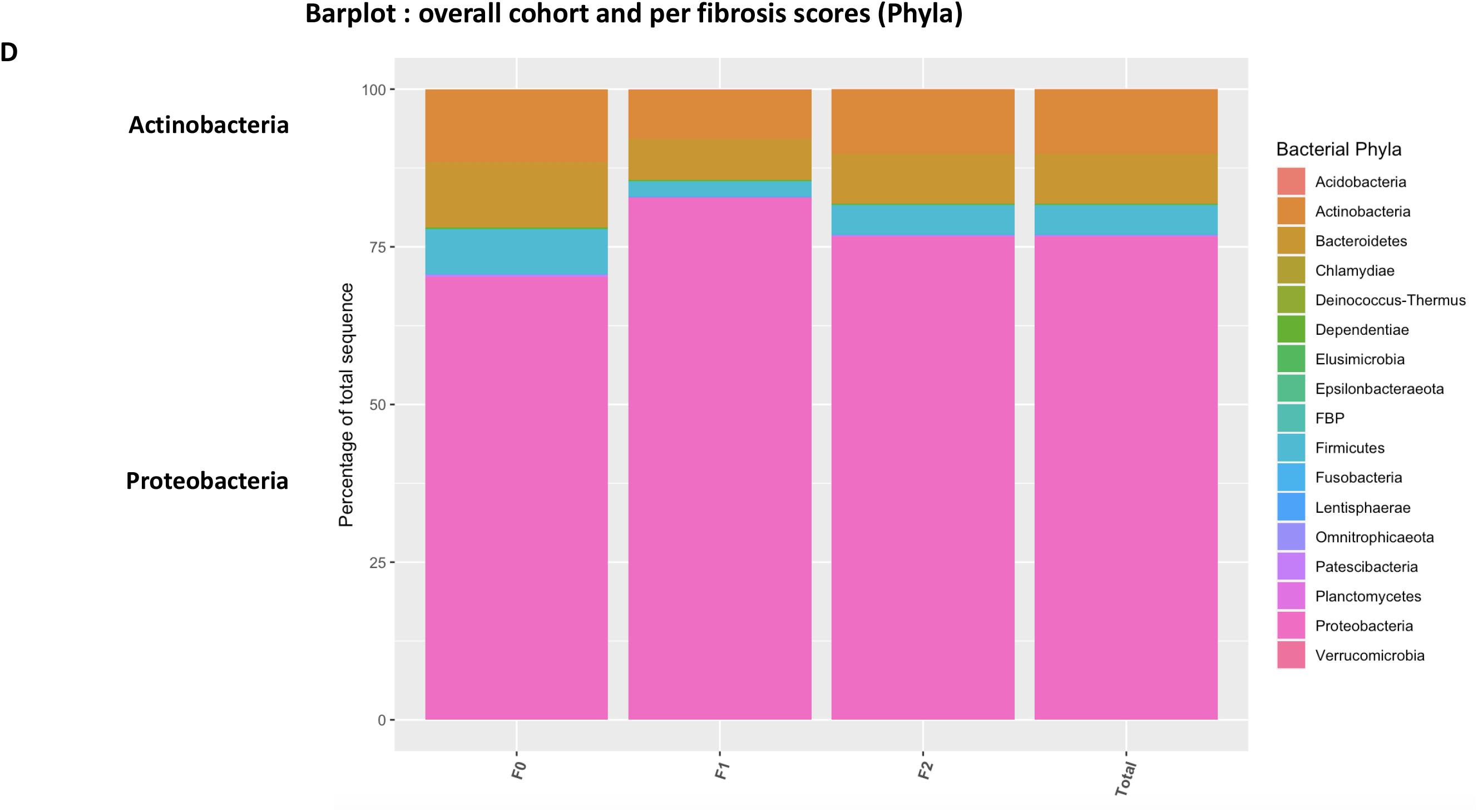
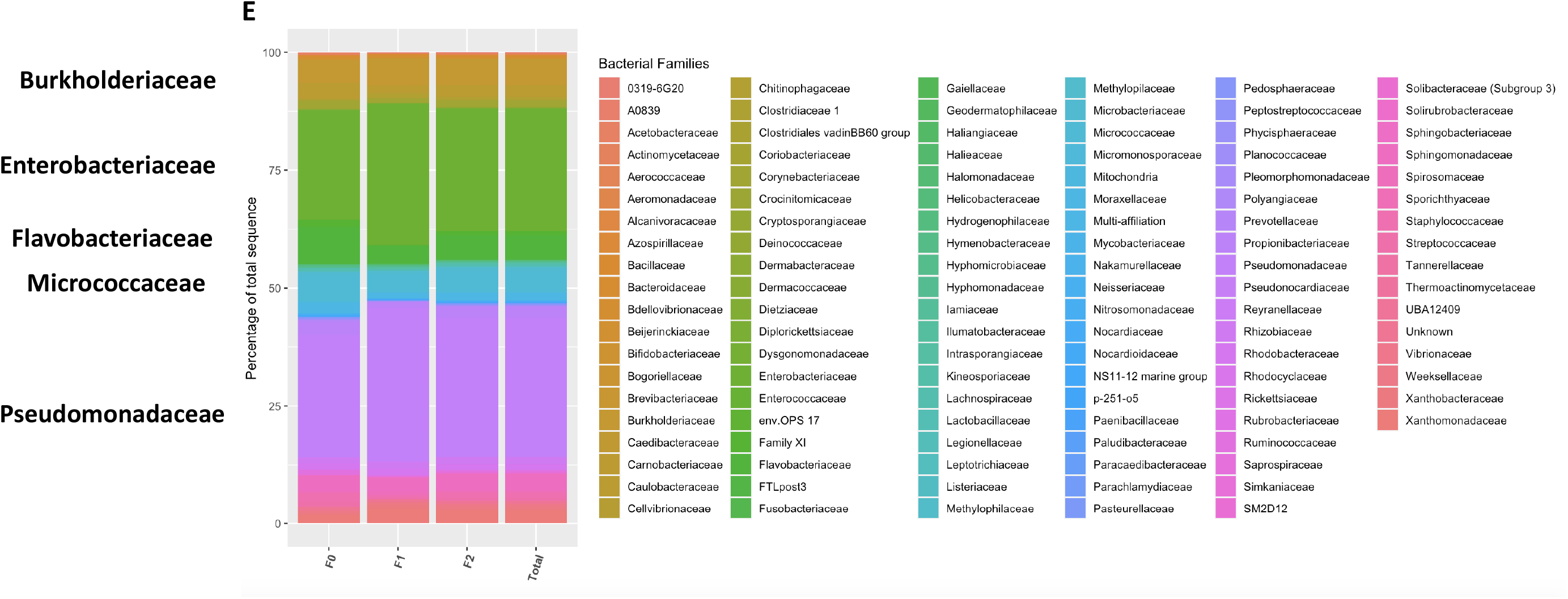
Visualization of liver 16SrDNA sequences by principal component analyses according to countries and fibrosis scores. The 16SrDNA OTUs sequences were used as entries for a principal component analysis (PCA). PCA-biplot from package Factoextra and FactomineR of individuals for the first two principal components are shown. They sum up 10.0% of the total variance of the dataset. Patients were grouped by **A**, countries (red dots=Austria, green triangle=Italy, blue square=Romania, purple cross=Spain) and by **B**, fibrosis scores (red dots=F0, green triangle=F1, blue square=F2). The vectors corresponding to the clinical variables are shown as arrows. **C** Barplot depicting the frequencies of liver microbial composition of each patient at the phylum level or **D** as means of the phyla frequencies or **E** the family frequencies for the overall cohort (total) or according to the fibrosis scores (F0, F1, F2).

To further identify whether liver fibrosis scores could be characterized by specific signatures we explored indexes of alpha and beta diversity of 16SrDNA in liver tissue. The data show that differences in abundances at the phylum, and family taxonomic levels were also associated with differences of the alpha diversity (**Supplementary Fig 2A,B,C)**. Notably, the Observed, Shannon and Simpson indexes were significantly different between the F0 and F1 groups at the phylum and family levels. In addition to alpha diversity, we analyzed beta diversity and performed a principal coordinate analysis (PCoA) considering distances between variables (using Bray-curtis distance). The PCoA analyses showed that the F0 group was distant from the two others which suggests a specific 16SrDNA signature (**Supplementary Fig 2D,E**). It is noteworthy that outlier patients were also detected. Although, when analyzed together the three groups could not be classified clearly. The F0 group differed graphically from the F1,F2 groups suggesting a specific signature discriminating between F0 and F1,F2. To determine if the ellipse centers of the F0 group differs from the ellipse center of the other groups, a Permutational Multivariate Analysis of variance (PERMANOVA) followed by a Kruskall-Wallis test were performed and found a difference between F0 and F1 groups (p<0.03). Along the same line of investigation, we performed different graphical representations such as heatmaps and Venn diagrams.

### Identification of specific bacterial signatures

To identify the variables that are specific to Fibrosis scores we performed a first Venn diagram on the overall set of variables (**Fig 3A**). Eighty-nine variables were common to all groups and considered as the core of the cohort while 21, 77, and 108 OTUs were specific of the F2, F1, F0 groups, respectively. To isolate extremely rare variables and unbalanced distribution between groups we next considered only OTUs with more than 25% of non-zero counts and an average number of counts per group higher than 150 and similarly drew a second Venn diagram. We identified 12, 5, and 5 OTUs specific to F2, F1, and F0 scores, respectively (**Fig 3B**) and (**Table 2**). To identify if these specific OTUs could be picked up using another approach we generated a heatmap where each OTUs was positioned while the fibrosis scores was fixed (**Fig 3C**). We noted that the frequencies of the majority of OTUs equal 0 or are extremely low (<0.01%) thereby, most of these variables do not bring information. Similarly, a minority of the variables of high frequencies were common to all liver fibrosis groups and did not provide discriminant information neither. Such OTUs could be considered as the core variable of liver microbiota. Conversely, a subset of OTUs could be considered as discriminant that was identified on a different heatmap following the removing of the non-informative OTUs (**Fig 3D**).

**TABLE 2:** Identification of specific bacterial signatures (unfair analyses).

| **F0** |  |  |  |
| --- | --- | --- | --- |
| **OTU name** | **Phylum** | **Family** | **Genus** |
| Cluster_30 | Proteobacteria | Moraxellaceae | Acinetobacter |
| Cluster_74 | Firmicutes | Ruminococcaceae | Faecalibacterium |
| Cluster_35 | Actinobacteria | Microbacteriaceae | Rhodoluna |
| Cluster_28 | Actinobacteria | Micrococcaceae | Kocuria |
| Cluster_21 | Proteobacteria | Caulobacteraceae | Caulobacter |

| **F1** |  |  |  |
| --- | --- | --- | --- |
| **OTU name** | **Phylum** | **Family** | **Genus** |
| Cluster_43 | Proteobacteria | Pseudomonadaceae | Pseudomonas |
| Cluster_31 | Proteobacteria | Pseudomonadaceae | Pseudomonas |
| Cluster_25 | Proteobacteria | Enterobacteriaceae | Multi-affiliation |
| Cluster_37 | Proteobacteria | Rhodobacteraceae | Paracoccus |
| Cluster_40 | Bacteroidetes | Chitinophagaceae | Ferruginibacter |

| **F2** |  |  |  |
| --- | --- | --- | --- |
| **OTU name** | **Phylum** | **Family** | **Genus** |
| Cluster_122 | Firmicutes | Lachnospiraceae | Multi-affiliation |
| Cluster_59 | Actinobacteria | Corynebacteriaceae | Corynebacterium<br>1 |
| Cluster_34 | Bacteroidetes | Weeksellaceae | Cloacibacterium |
| Cluster_48 | Firmicutes | Peptostreptococcaceae | Romboutsia |
| Cluster_42 | Proteobacteria | Burkholderiaceae | Tepidimonas |
| Cluster_26 | Firmicutes | Streptococcaceae | Lactococcus |
| Cluster_53 | Bacteroidetes | Weeksellaceae | Cloacibacterium |
| Cluster_18 | Proteobacteria | Enterobacteriaceae | Pantoea |
| Cluster_115 | Proteobacteria | Burkholderiaceae | Delftia |
| Cluster_50 | Actinobacteria | Microbacteriaceae | Clavibacter |
| Cluster_91 | Actinobacteria | Corynebacteriaceae | Corynebacterium |
| Cluster_54 | Proteobacteria | Burkholderiaceae | Ralstonia |

Clusters were identified from the overall database prior to applying the fair strategies. The impact of countries is observed as shown in Fig 2.

**Fig. 3:**
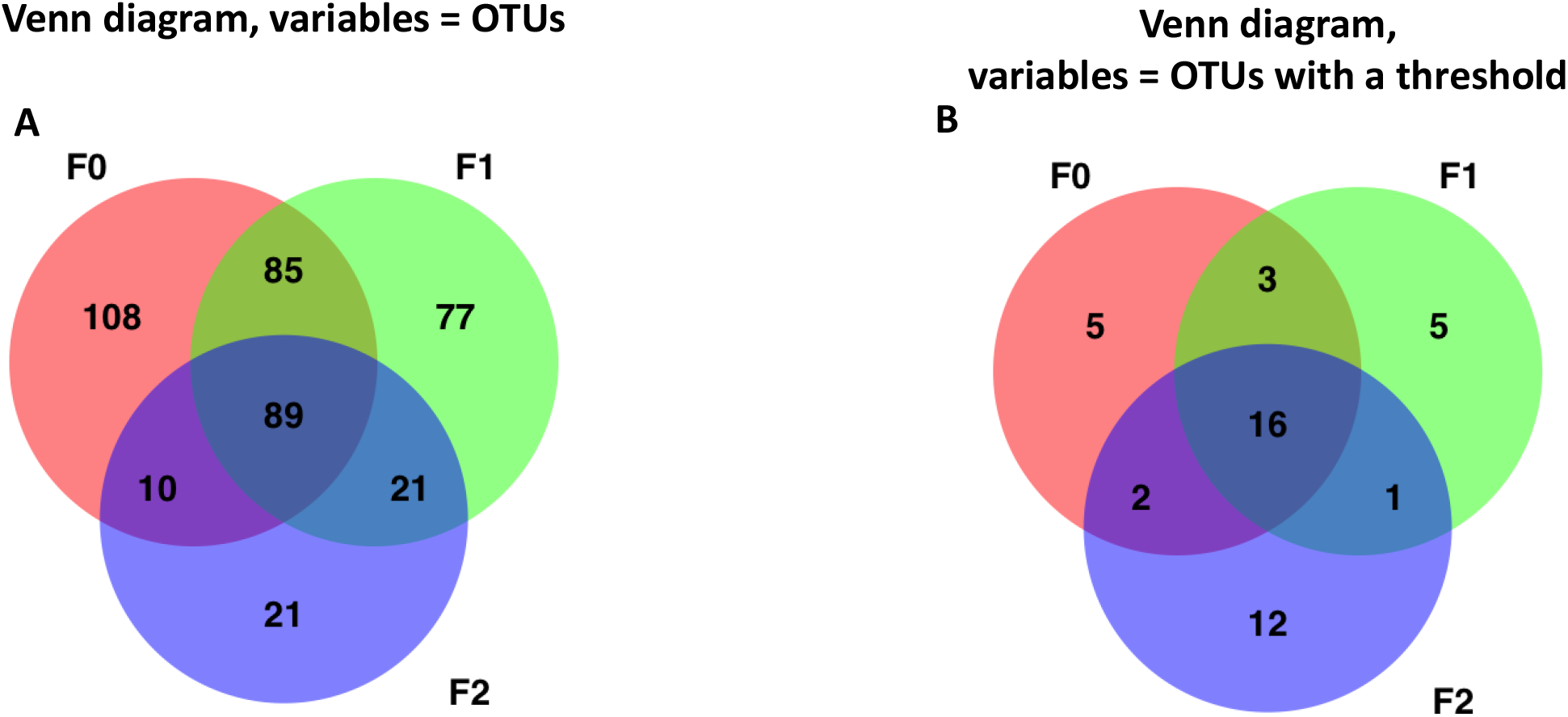
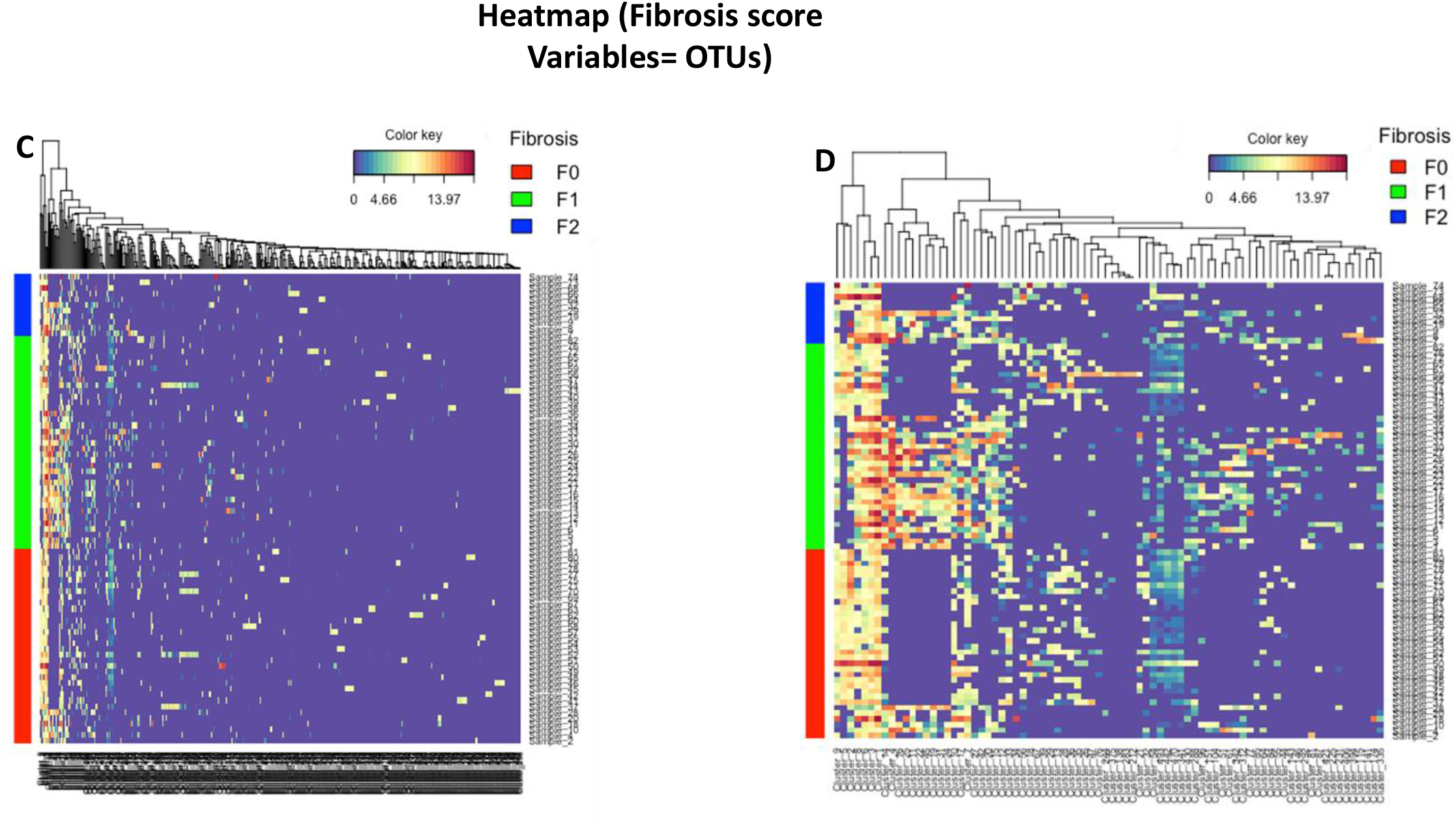
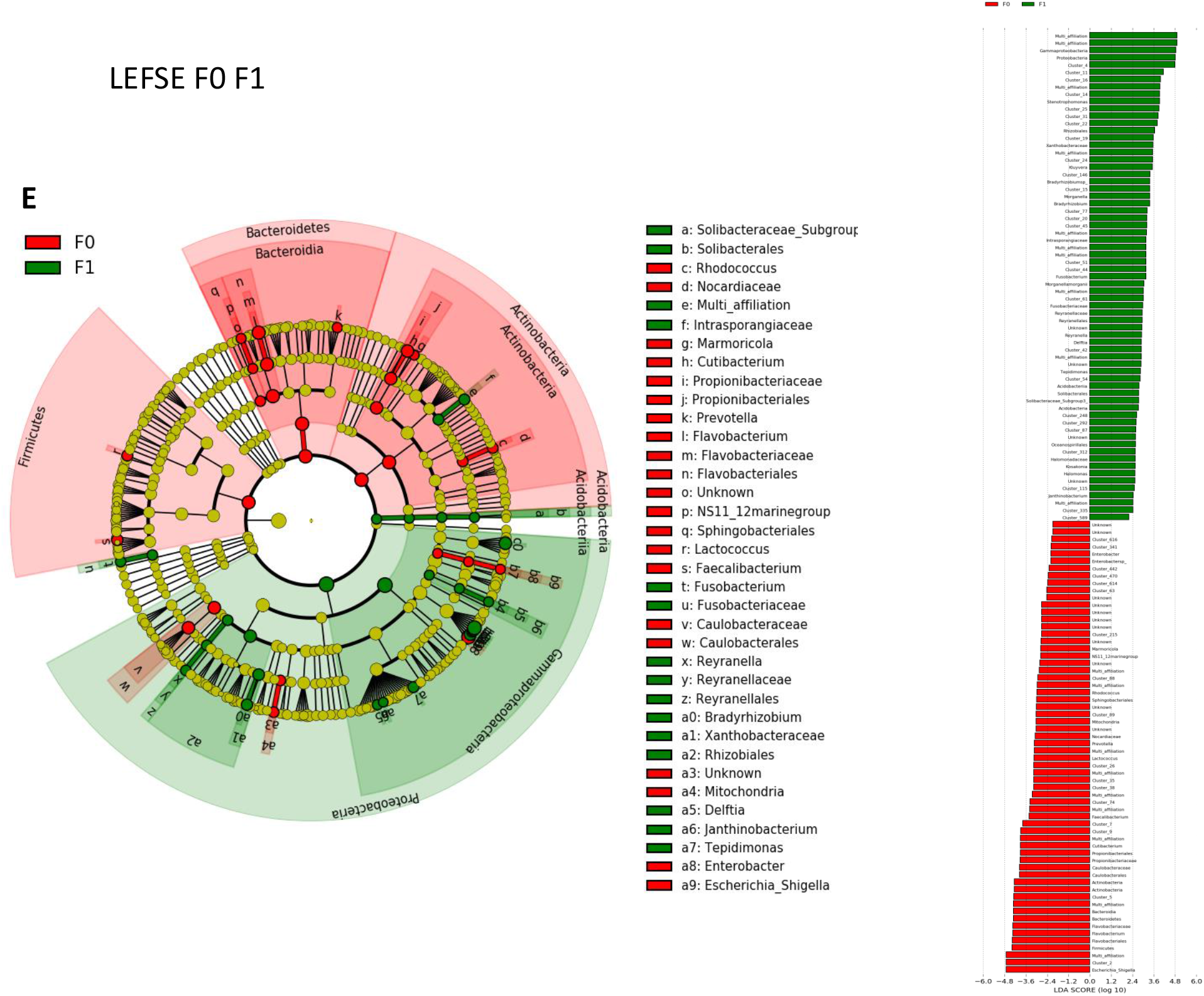
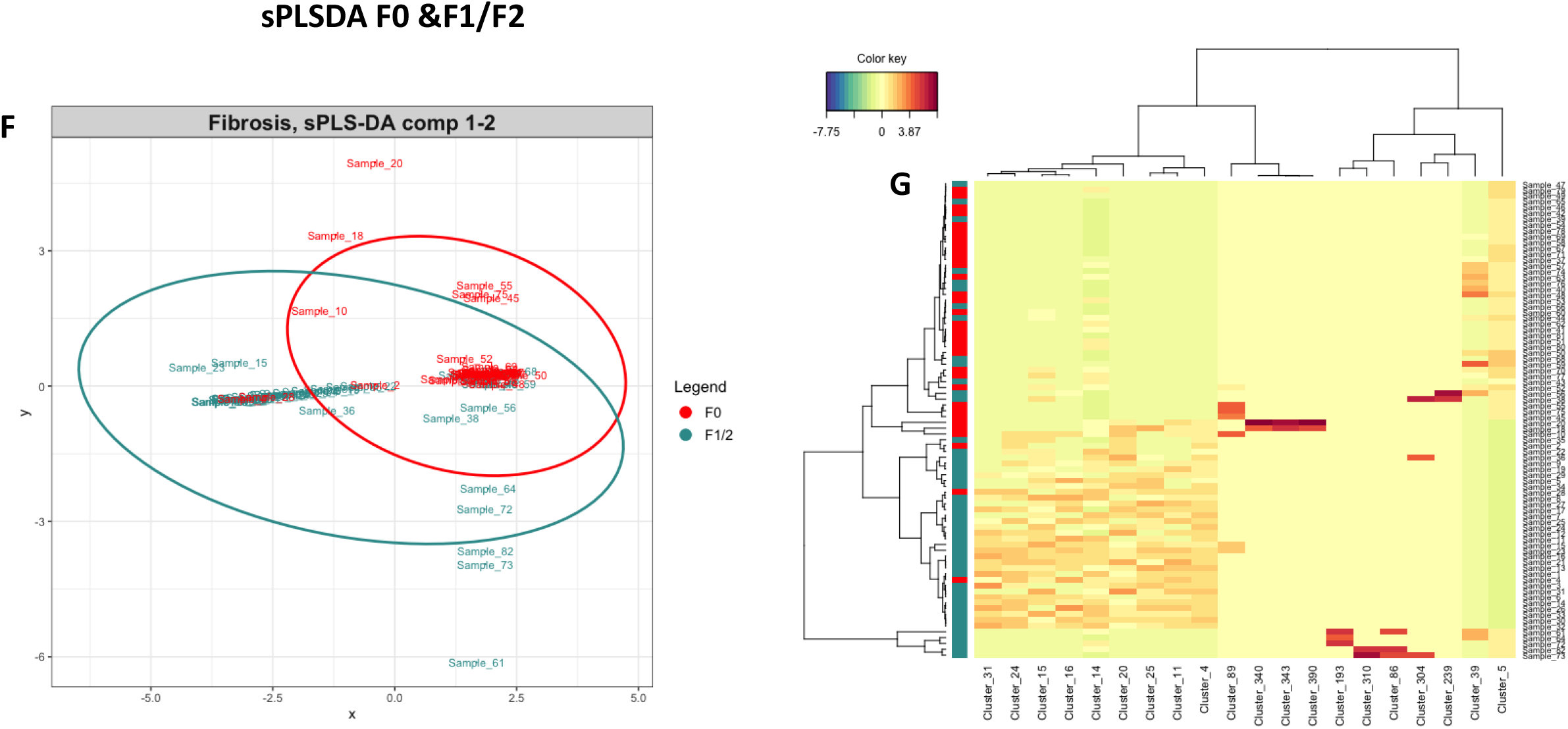
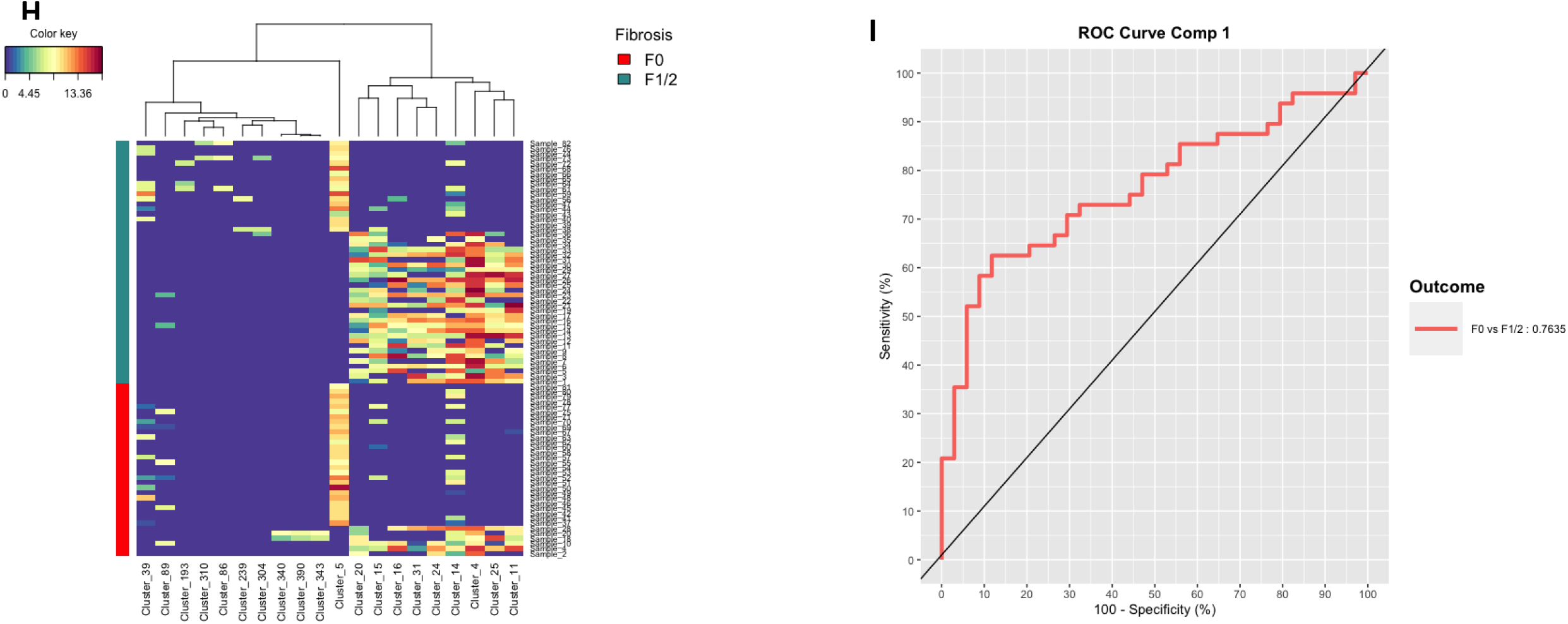
Discriminant analysis strategies of the liver microbiota 16SrDNA OTUs according to the fibrosis scores. Venn diagrams where **A** all the 16SrDNA taxa or **B** data after removing those extremely rare and with unbalanced distribution within the 3 groups of patients with liver fibrosis, were used as entry variables characterizing the 3 liver fibrosis scores (red=F0, green=F1, blue=F2). **C** Heatmap of normalized OTU counts according to the 3 groups of patients with liver fibrosis scores and **D** a corresponding subset of normalized OTU counts with groups of patients fixed. **E** LEfSe cladogram of taxonomic assignments from 16SrDNA sequence data of the two liver biopsy fibrosis groups (F0 and F1). The cladogram shows the taxonomic levels represented by rings with phyla at the innermost ring and genera at the outermost ring, and each circle is a member within that level. Taxa at each level are shaded according to the liver fibrosis group in which it is more abundant (P < 0.05; LDA score ≥2.0). LDA scores are shown on the right panel for each taxon. **F** sPLSDA classification performance on a CSS normalized microbial table count of the F0 versus F1/2 groups of patients. Sample plot, each point corresponds to an individual and is colored according to its fibrosis score (red=F0, green=F1/2). **G** Clustering Image Map (CIM) of the OTUs selected on each sPLS-DA component. **H** Heatmap of the OTUs selected on each sPLS-DA component with groups of patients fixed. **I** ROC calculated on the predicted scores obtained from the sPLSDA model.

To refine the identifications of the discriminant bacteria we performed a Linear discriminant analysis (LDA) coupled with effect size measurements (**Fig 3E, Supplementary Fig 3A,B**). The data show that most of the discriminant information was identified when comparing between F0 and F1. The Firmicutes, Flavobacteriaceae, Caulobacteraceae and Actinobacteria were specific to F0 group and the Proteobacteria was specific to F1 (**Fig 3F**). On the boxplot the taxa enriched in patients with no fibrosis are indicated with a negative score and mild fibrosis enriched taxa are indicated with a positive score. We performed LEFSe between each score pairs and identified much less differences between F1 F2 suggesting that they could have a similar liver microbiota, as suggested in **Fig 2B** despite the discriminant clinical variables identified in **Fig1B**.

On these first sets of analyses, the number of fibrosis scores of each patient was too heterogeneous to perform a discriminant analysis (overfitting). As shown on **supplementary Fig 2D**,**3A** they were almost no difference between F1 and F2, therefore we merged F1 and F2 scores as F1/2 group, increasing hence the number of patients of that group.

To validate the pertinence of such strategy we performed a partial least square discriminant analysis i.e. PLS-DA. To select the most discriminative features in the model we used its sparse version sPLS-DA based on a Lasso penalization. The number of variables to be selected per component involved in the visualization is optimized using leave-one-out cross-validation. On the sample plot (**Fig 3F**), we observe a slight separation of the two fibrosis scores ellipses compared to the unsupervised PCA. From the most discriminant OTUs selected on each sPLS-DA component, a dissociation between the two groups can be visualized using a Clustering Image Map (CIM) (**Fig 3 G,H**). The graphs show a clear classification of the patients based on the identified discriminant variables. Eventually, we calculated the ROC curve with all discriminant variables that shows an increased specificity and sensitivity above baseline (**Fig 3I**).

Altogether, some degree of graphical classification of the liver fibrosis score could be observed using the clinical database and the 16SrDNA database. However, in both instances the individuals appear to be still distributed according to the countries. Therefore, to overcome this issue we developed an ad hoc fairness statistical strategy allowing the classification of variables i.e. OTUs independently from the cohort.

### Identification of clusters of cohort-independent 16SrDNA associated with different mild scores of fibrosis

In front of these numerous signatures and the influence of confounding factors such as the impact of the cohort set there is a need to identify clusters of variables specific to each liver fibrosis score but independent of the cohort impact. To this aim we considered three different fair approaches on the overall cohorts and then defined clusters of OTU variables independent from the cohort. The first fair approach consists in identifying principal components from the metagenomic dataset as signatures of the cohorts and removing them to generate a new dataset where no components would be cohort sensitive. To this aim we compared the largest cohort i.e. from Romania to the others. Principal components conditional distributions with respect to the cohorts were visualized (**Fig 4A**). Then, we removed the principal components the most correlated with the cohorts when the absolute value of Pearson correlation was above a threshold. The remaining non-overlapping components are cohort-insensitive and used to identify the variables associated with the mild fibrosis score. Remarkably, more than 78% of the variation from the original data was still included into the selected principal components suggesting that the discriminant information was only marginally affecting our previous results. On this “fair” dataset we applied the standard random forest classification to predict fibrosis scores. From the variable importance plot, indicating the contribution of the variables to classify the data, we selected the 10 more predictive principal components and identified 3 significantly associated with fibrosis scores (**Fig 4B,C,D**).

**Fig. 4:**
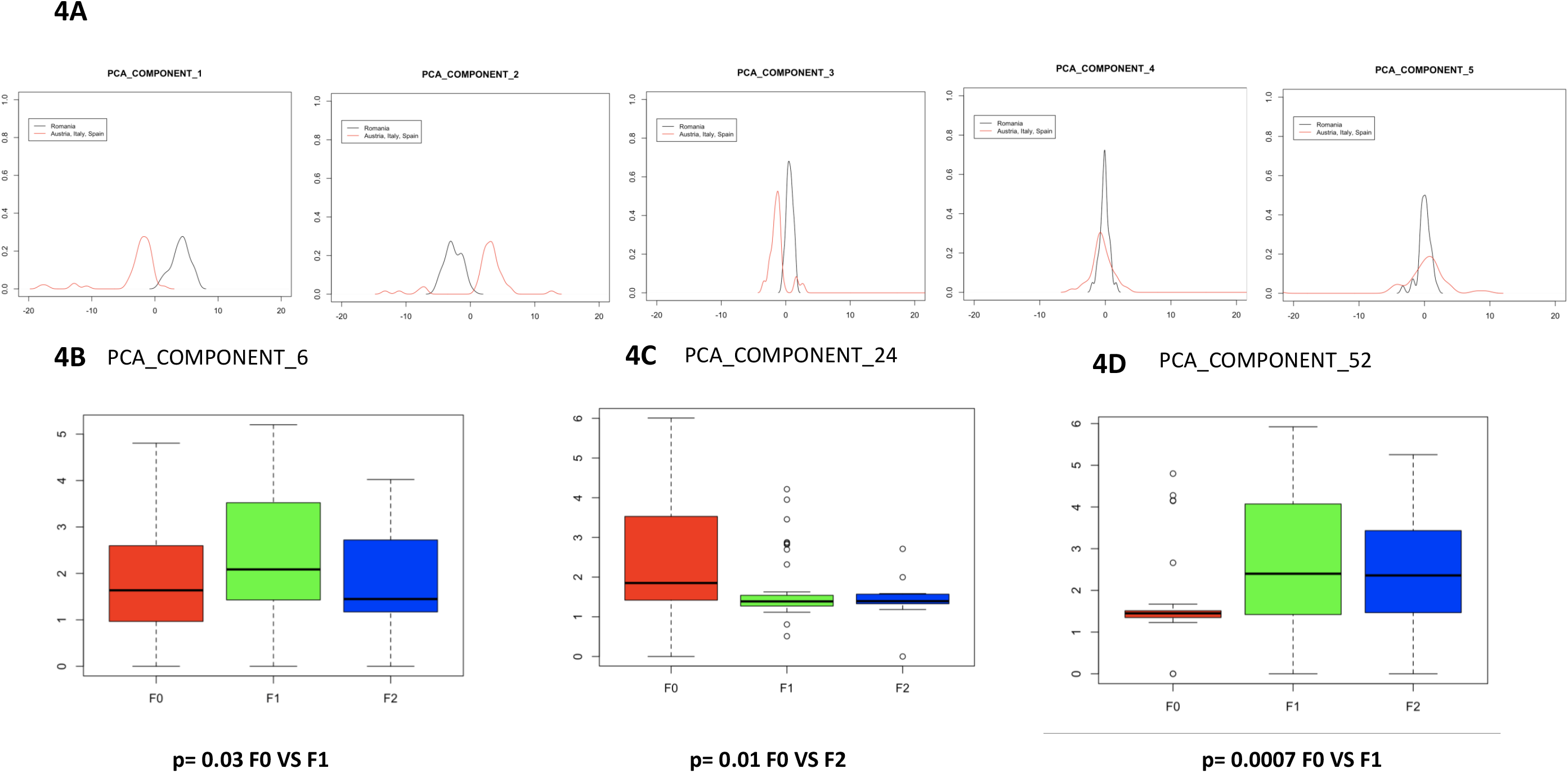
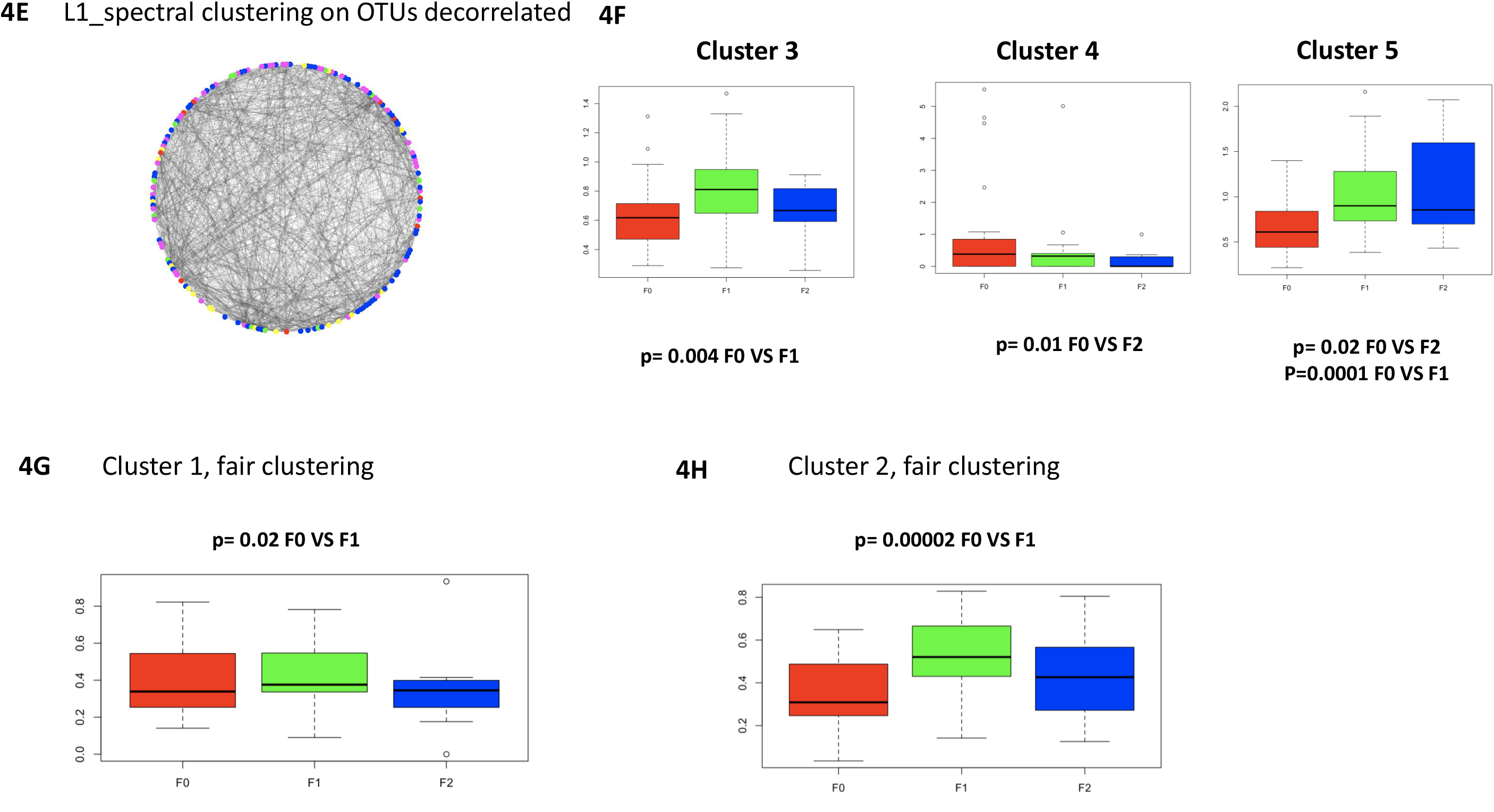
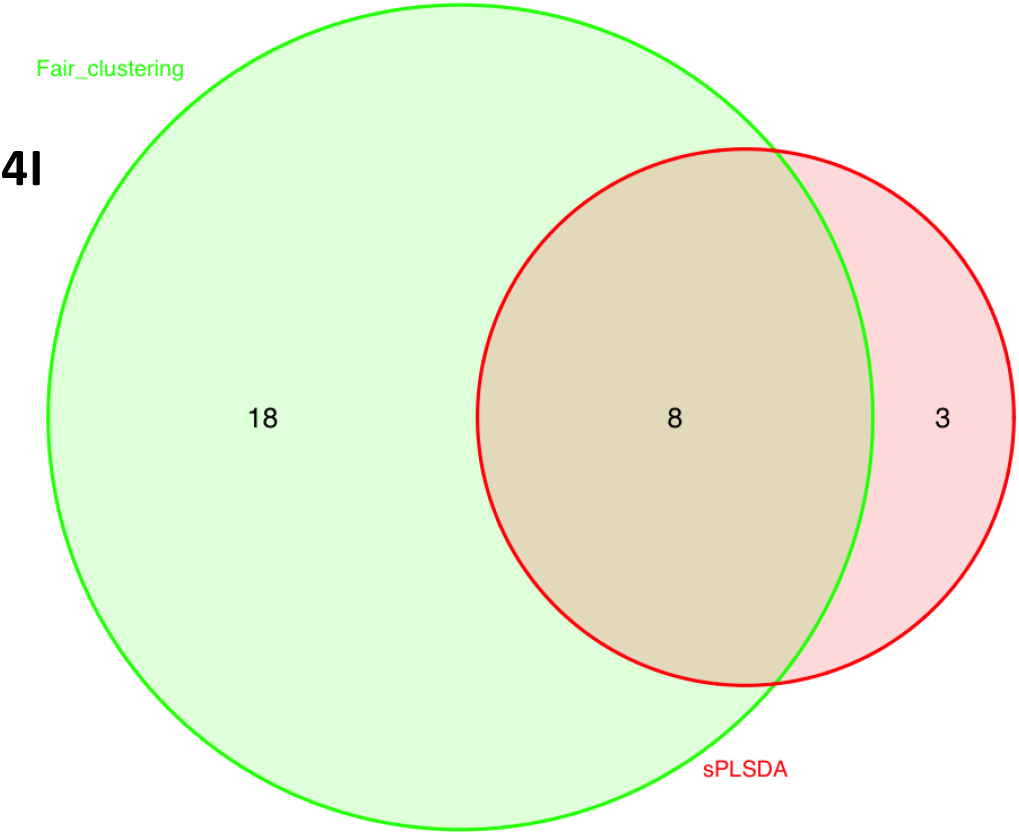
discriminant analyses of the 16SrDNA OTUs variables using fairness strategies. **A** Distribution curves (or densities) of the coordinate of individuals, split into two cohort types (black=Romania, red= the other countries: Italy, Austria, and Spain), when projected on the five first principal components built from the 16SrDNA OTUs normalized table count. The non-overlapping plots (for example components 1,2,3) correspond to cohort discriminant components and will be removed from the final analysis to identify the liver fibrosis discriminant variables. Boxplot representing the frequencies of the most significant OTUs contributing to **B** the 6^th^, **C** the 24^th^, **D** the 52^nd^ principal components for the different groups of liver fibrosis scores (red=F0, green=F1, blue=F2). **E** Graphical representation of the normalized OTU table counts whose nodes are colored according to the 5 clusters identified by the l1-spectral clustering algorithm (red= 1, green= 2, blue, 3, pink= 4 and yellow= 5). **F** Boxplot representing the mean frequencies of the OTUs in cluster 3, 4 and 5, identified by the l1-spectral clustering algorithm, for the different groups of liver fibrosis scores (red=F0, green=F1, blue=F2). **G, H** Boxplot representing the frequencies of OTUs in cluster 1, and 2, identified by fair-tree algorithm, for the different groups of liver fibrosis scores (red=F0, green=F1, blue=F2). **I** Venn diagram depicting the liver microbial taxonomies of common OTUs identified by standard (sPLS-DA) and fair approaches (fairtree, random forest, l1-spectral clustering) as signatures of low fibrosis scores (green= sPLSDA, red= fair algorithms).

The second fair clustering approach directly selects OTUs which are the most influenced by the cohorts and removes them from the analysis. The associated matrix is then modeled by a graph and subjected to a spectral clustering algorithm to which we applied an L1 penalty. The nodes represent OTUs and the edges show interactions between each pair of variables (**Fig 4E**). Using this novel l1-spectral clustering algorithm we identified 5 clusters of OTUs among which 3 were significantly associated with the liver fibrosis scores (**Fig 4F**).

Eventually, we performed the fair clustering method called “fair-tree”. We used the 16SrDNA normalized table count to identify clusters with approximately equal number of patients from each cohort. Two of the three clusters found containing respectively 36 and 97 OTUs, were statistically significant when comparing F0 versus F1 scores (**Fig 4G,H**).

To summarize all the identified OTUs significantly associated with the different low scores of fibrosis, we compiled them in (**Table 3**) and identified their respective taxa. From the fair principal components identified, we only considered the five OTUs that contribute most to create each of these components. Then, from the Venn diagram we identified common OTUs signatures of low fibrosis scores from standard (sPLS-DA) and fair approaches (fair-tree, random forest, l1-spectral clustering) (**Fig 4I**). Interestingly, from all selected OTUs eight common OTUs were from the same phylum i.e. Proteobacteria (**Table 4**) suggesting that most of the discriminant information could be due to these taxa. However, there is still most likely some information that this predominant family could be hiding. We therefore set a new mathematical strategy to exemplify the low frequency and meaningful bacteria by using the TF-IDF (Term frequency-inverse document frequency) approach.

**TABLE 3.** Identification of clusters of cohort-independent 16SrDNA associated with the different low scores of fibrosis.

| **sPLSDA** |  |  |  |  |
| --- | --- | --- | --- | --- |
| **OTU name** | **Phylum** | **Family** | **Genus** | **Significance** |
| Cluster_20 | Proteobacteria | Burkholderiaceae | Ralstonia | F0 VS F1 |
| Cluster_15 | Proteobacteria | Xanthobacteraceae | Bradyrhizobium | F0 VS F1 |
| Cluster_16 | Proteobacteria | Enterobacteriaceae | Multi-affiliation | F0 VS F1 |
| Cluster_31 | Proteobacteria | Pseudomonadaceae | Pseudomonas | F0 VS F1 |
| Cluster_24 | Proteobacteria | Enterobacteriaceae | Kluyvera | F0 VS F1 |
| Cluster_14 | Proteobacteria | Xanthomonadaceae | Stenotrophomonas | F0 VS F1 |
| Cluster_4 | Proteobacteria | Enterobacteriaceae | Multi-affiliation | F0 VS F1 |
| Cluster_25 | Proteobacteria | Enterobacteriaceae | Multi-affiliation | F0 VS F1 and F0 VS F2 |
| Cluster_11 | Proteobacteria | Pseudomonadaceae | Pseudomonas | F0 VS F1 |
| Cluster_89 | Actinobacteria | Corynebacteriaceae | Corynebacterium 1 | F0 VS F1 and F0 VS F2 |
| Cluster_5 | Bacteroidetes | Flavobacteriaceae | Flavobacterium | F0 VS F1 and F0 VS F2 |

| **Fair-tree** |  |  |  |  |
| --- | --- | --- | --- | --- |
| **OTU name** | **Phylum** | **Family** | **Genus** | **Significance** |
| Cluster_31 | Proteobacteria | Pseudomonadaceae | Pseudomonas | F0 VS F1 |
| Cluster_66 | Proteobacteria | Pasteurellaceae | Haemophilus | F1 VS F2 and F0 VS F2 |
| Cluster_335 | Proteobacteria | Burkholderiaceae | Janthinobacterium | F0 VS F1 |
| Cluster_42 | Proteobacteria | Burkholderiaceae | Tepidimonas | F0 VS F1 |
| Cluster_341 | Proteobacteria | Enterobacteriaceae | Enterobacter | F0 VS F1 |
| Cluster_248 | Proteobacteria | Reyranellaceae | Reyranella | F0 VS F1 and F0 VS F2 |
| Cluster_231 | Proteobacteria | Enterobacteriaceae | Multi-affiliation | F0 VS F1 |
| Cluster_36 | Bacteroidetes | Flavobacteriaceae | Flavobacterium | F1 VS F2 and F0 VS F2 |
| Cluster_64 | Actinobacteria | Corynebacteriaceae | Lawsonella | F1 VS F2 and F0 VS F2 |
| Cluster_339 | Proteobacteria | Burkholderiaceae | Limnohabitans | F1 VS F2 |
| Cluster_15 | Proteobacteria | Xanthobacteraceae | Bradyrhizobium | F0 VS F1 |
| Cluster_44 | Actinobacteria | Intrasporangiaceae | Multi-affiliation | F0 VS F1 and F0 VS F2 |
| Cluster_25 | Proteobacteria | Enterobacteriaceae | Multi-affiliation | F0 VS F1 |
| Cluster_45 | Proteobacteria | Burkholderiaceae | Comamonas | F0 VS F1 |
| Cluster_14 | Proteobacteria | Xanthomonadaceae | Stenotrophomonas | F0 VS F1 |

| **Fair Random Forest** |  |  |  |  |
| --- | --- | --- | --- | --- |
| **OTU name** | **Phylum** | **Family** | **Genus** | **Significance** |
| Cluster_14 | Proteobacteria | Xanthomonadaceae | Stenotrophomonas | F0 VS F1 |
| Cluster_77 | Proteobacteria | Enterobacteriaceae | Multi-affiliation | F1 VS F2 and F0 VS F2 |
| Cluster_35 | Actinobacteria | Microbacteriaceae | Rhodoluna | F0 VS F1 |
| Cluster_36 | Bacteroidetes | Flavobacteriaceae | Flavobacterium | F0 VS F1 |
| Cluster_16 | Proteobacteria | Enterobacteriaceae | Multi-affiliation | F0 VS F1 and F0 VS F2 |
| Cluster_312 | Proteobacteria | Enterobacteriaceae | Kosakonia | F0 VS F1 |
| Cluster_24 | Proteobacteria | Enterobacteriaceae | Kluyvera | F0 VS F1 |

| Fair l1_spectral clustering |  |  |  |  |
| --- | --- | --- | --- | --- |
| OTU name | Phylum | Family | Genus | Significance |
| Cluster_31 | Proteobacteria | Pseudomonadaceae | Pseudomonas | F0 VS F1 |
| Cluster_15 | Proteobacteria | Xanthobacteraceae | Bradyrhizobium | F0 VS F1 |
| Cluster_341 | Proteobacteria | Enterobacteriaceae | Enterobacter | F0 VS F1 |
| Cluster_20 | Proteobacteria | Burkholderiaceae | Ralstonia | F0 VS F1 |
| Cluster_4 | Proteobacteria | Enterobacteriaceae | Multi-affiliation | F0 VS F1 |
| Cluster_2 | Proteobacteria | Enterobacteriaceae | Escherichia-Shigella | F0 VS F1 |
| Cluster_27 | Proteobacteria | Enterobacteriaceae | Serratia | F0 VS F1 |
| Cluster_3 | Proteobacteria | Pseudomonadaceae | Pseudomonas | F0 VS F1 |
| Cluster_24 | Proteobacteria | Enterobacteriaceae | Kluyvera | F0 VS F1 |
| Cluster_22 | Proteobacteria | Burkholderiaceae | Multi-affiliation | F0 VS F1 |
| Cluster_16 | Proteobacteria | Enterobacteriaceae | Multi-affiliation | F0 VS F1 |

**TABLE 4.** Microbial signatures common to all strategies.

| OTU name | Phylum | Family | Genus | Species | Significance |
| --- | --- | --- | --- | --- | --- |
| Cluster_31 | Proteobacteria | Pseudomonadaceae | Pseudomonas | Pseudomonas putida | F0 VS F1 |
| Cluster_15 | Proteobacteria | Xanthobacteraceae | Bradyrhizobium | Bradyrhizobium sp. | F0 VS F1 |
| Cluster_25 | Proteobacteria | Enterobacteriaceae | Multi-affiliation | Multi-affiliation | F0 VS F1 |
| Cluster_14 | Proteobacteria | Xanthomonadaceae | Stenotrophomonas | Multi-affiliation | F0 VS F1 |
| Cluster_16 | Proteobacteria | Enterobacteriaceae | Multi-affiliation | Multi-affiliation | F0 VS F1 |
| Cluster_24 | Proteobacteria | Enterobacteriaceae | Kluyvera | Multi-affiliation | F0 VS F1 |
| Cluster_20 | Proteobacteria | Burkholderiaceae | Ralstonia | Multi-affiliation | F0 VS F1 |
| Cluster_4 | Proteobacteria | Enterobacteriaceae | Multi-affiliation | Multi-affiliation | F0 VS F1 |

### Low frequency bacterial 16SrDNA gene contains classifying information

From the table count of all significant OTUs detected we generated a “word-cloud” (**Fig 5A, B**) to visualize the most abundant TF-IDF transformed OTU counts, regardless of fibrosis scores when compared to those non-normalized. Cluster 2 emerged as the most important discriminant OTU (taxonomic identifaction=Bacteria|Proteobacteria|Gammaproteobacteria|Enterobacteriales|Enterobacteriaceae|Escherichia-Shigella) further confirming the important amount of information contained in the Proteobacteria.

**Fig. 5:**
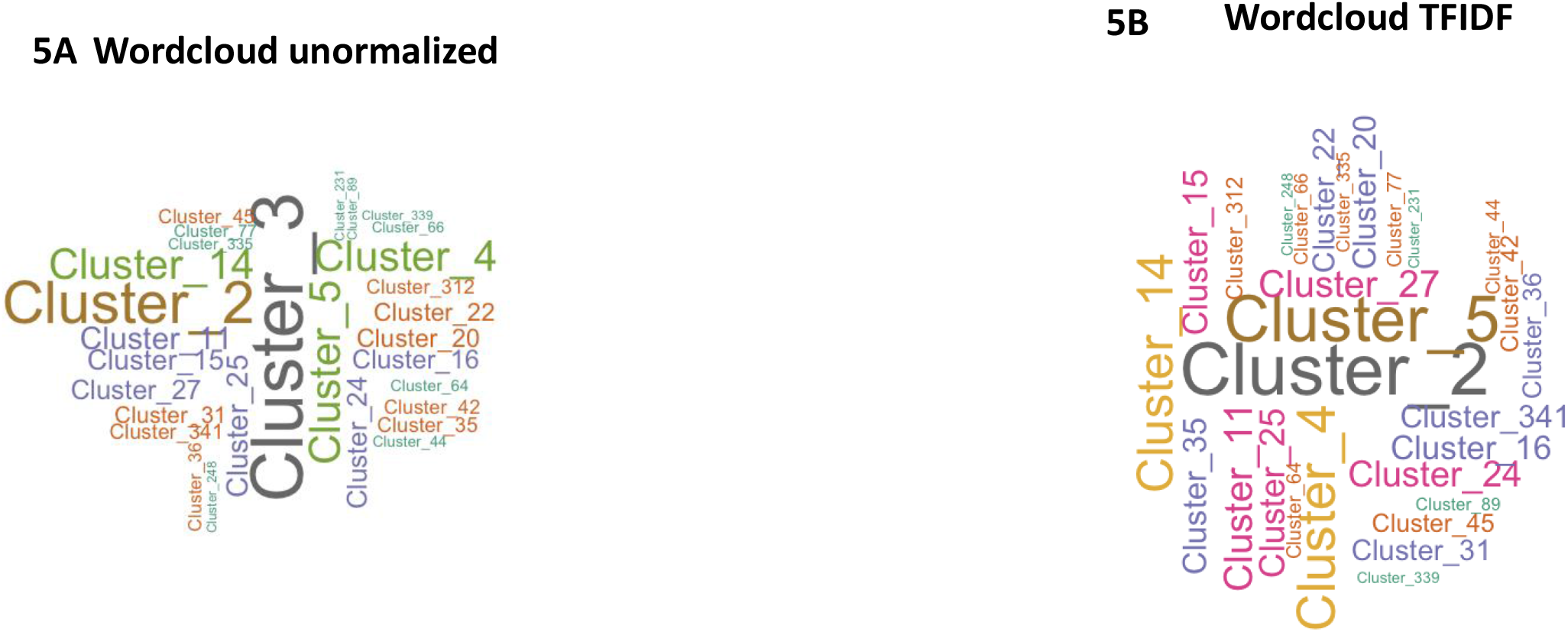
identification of clusters by wordclouds representation with or without TFIDF normalization. Wordclouds representing taxa of all significant bacteria according to **A**, their frequencies or **B**, after TFIDF normalization. The size of the name of bacteria is proportional to the frequency of the cluster in the cohorts.

Based on the identified specific signatures the next step was to generate hypotheses regarding their putative mode of action to the induction of the early events of liver fibrosis. We therefore performed predicted functional metagenomics.

### Predicted functional metagenome pathways

To identify the pathways and enzymes involved in the early development of fibrosis, we run predicted functional metagenomics algorithms based on the fairness-selected bacterial taxa. The heatmap shows clusters of enzymes that are associated with the F0 vs F1-2 fibrosis scores (**Fig 6A**). Eventually, sPLS-DA showed also a clear discrimination between the F0 vs F1-2 fibrosis scores. To evaluate the accuracy and sensitivity of our analyses as potential diagnostic tool, we drew a ROC and quantified the urea under curve with a score of 81.4% of accuracy (**Fig 6B,C**). We performed a similar analysis on pathways and showed specific clusters also discriminately associated with the fibrosis scores with a score of accuracy of 81.2% (ROC curve) (**Fig 6D-F**). We then represented an listed all selected enzymes and pathways highly expressed in the two major discriminant components (**Fig 6G-J**) and (**Table5**). Three pathways were highly and negatively associated with the liver fibrosis score of F1-2 when compared to the F0. We identified from the MetaCyc database (https://metacy.org/) that the preQ_0_ biosynthesis (PWY-6703), specific to Enterobacteriaceae such as E. coli, is involved notably in the synthesis of tetrahydrofolate and a class of nucleoside analogues that often possesses antibiotic, antineoplastic, or antiviral activities [29,30] (**Fig 6K**). In addition, two other pathways related to glucoryranose (PWY 6737) and glycogen (GLYCOCAT-PWY) degradation were identified probably providing energy to the main preQ_0_ biosynthesis pathway.

**Fig. 6:**
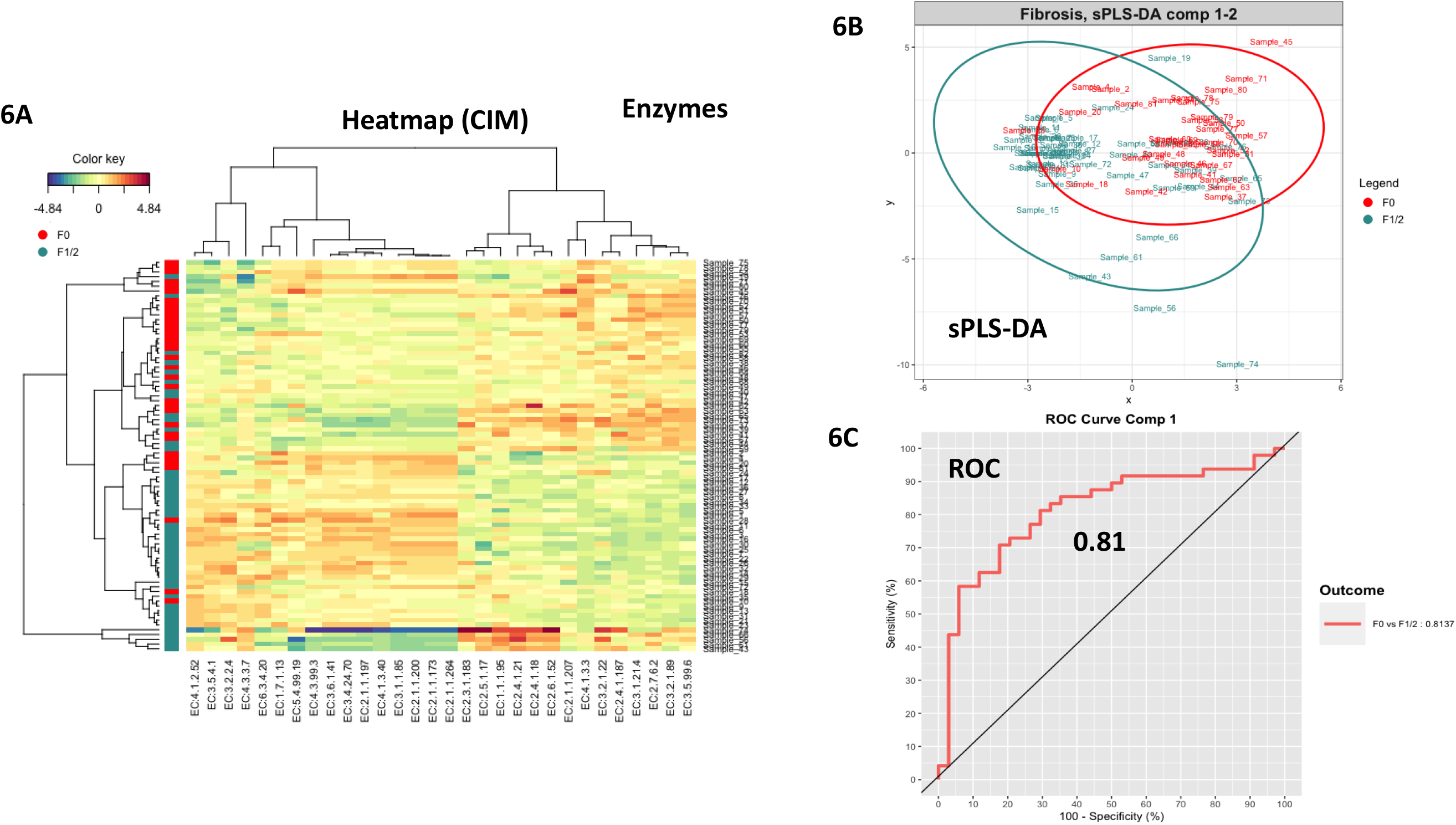
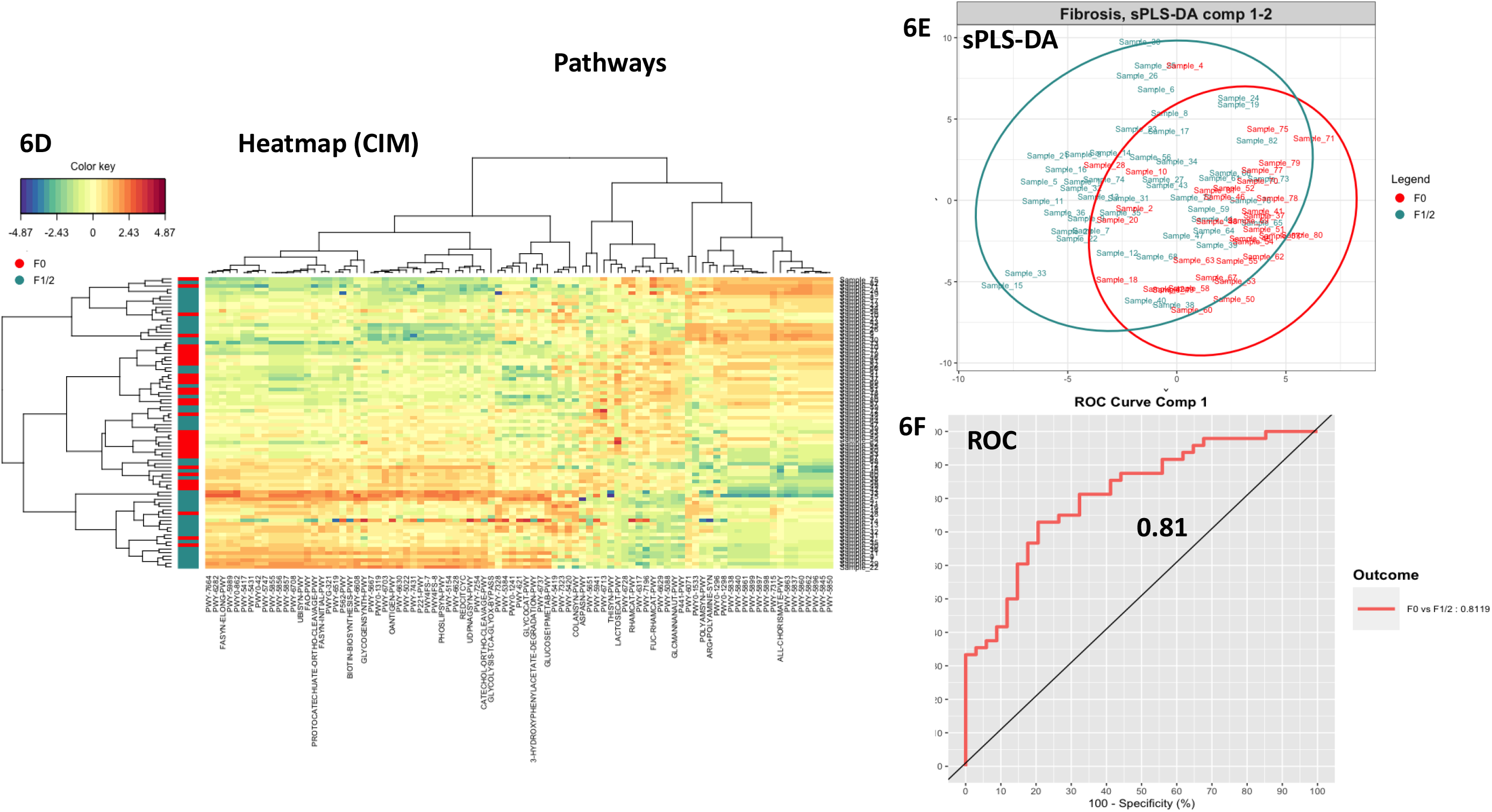
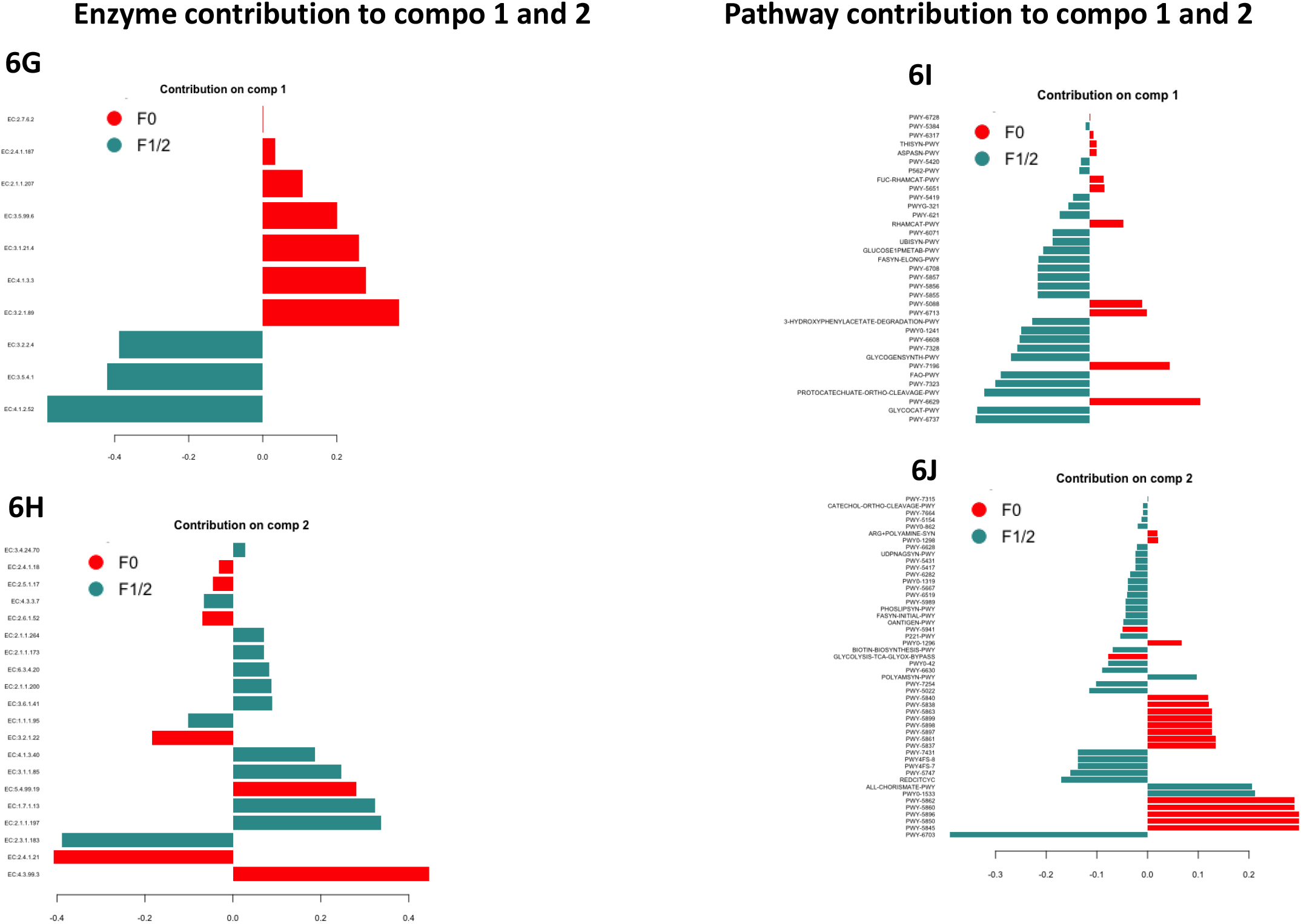
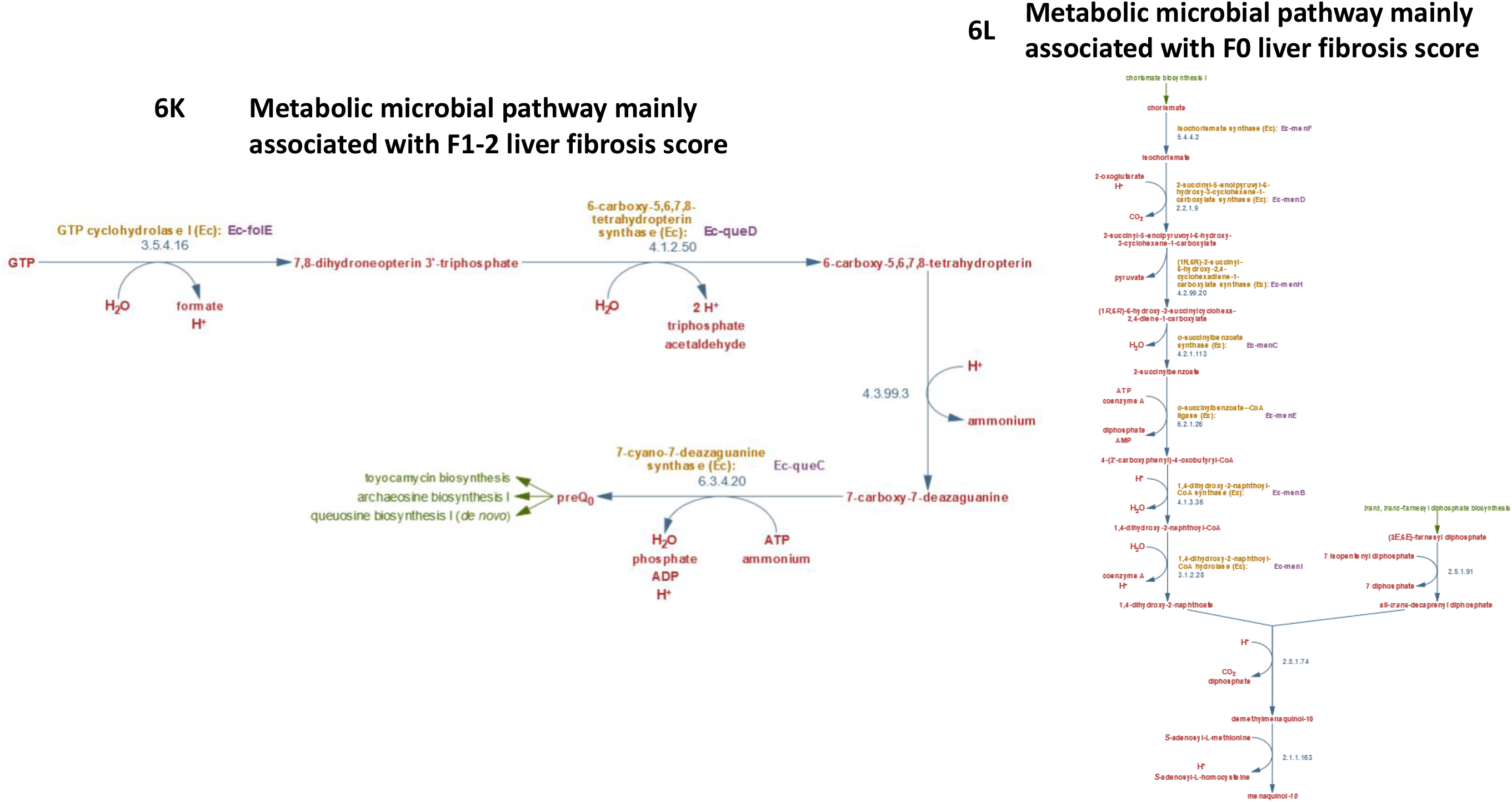
Predicted functional metagenomics analyses of discriminant enzymes and according to the fibrosis score. **A**,**D:** Heatmap (Clustering Image Map (CIM)), **B**,**E:** Sample plot, each point corresponds to an individual and is colored according to its liver fibrosis score (red=F0, green=F1/2), **C**,**F:** ROC classification performances of **A-C:** enzymes, and **D-F:** pathways, on a CSS normalized enzyme table count of the F0 versus F1/2 groups of patients. **G-I:** Loading plot representing the contribution of each enzyme (**G**,**H**), and pathways (**I**,**J**) selected to build the first and second components (red=F0, green=F1/2). **K**,**L:** main metabolic pathways from the MetaCyc database identified from the Loading plots for the **K:** F1-2 and **K:** F0 liver fibrosis scores.

**Table 5:**
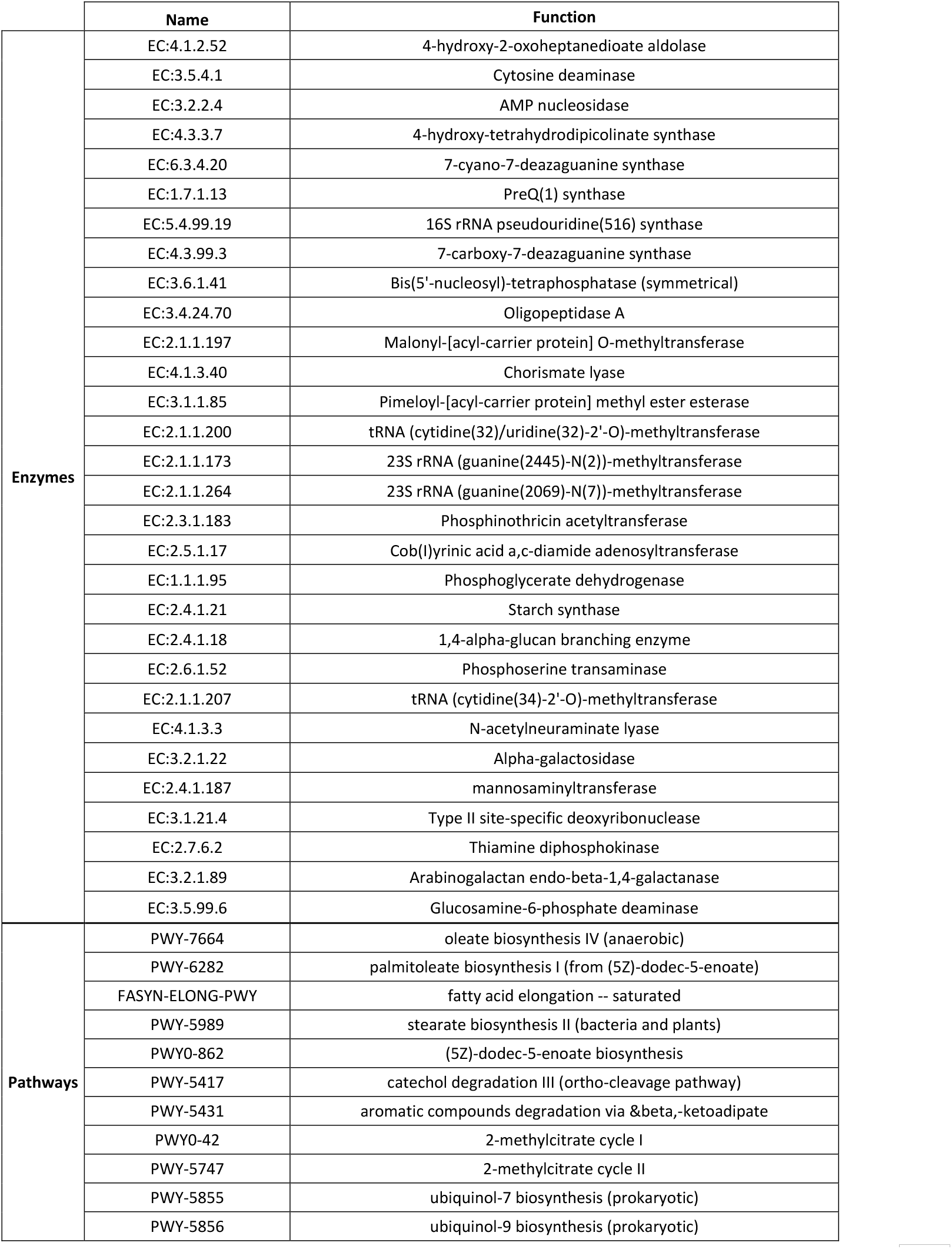
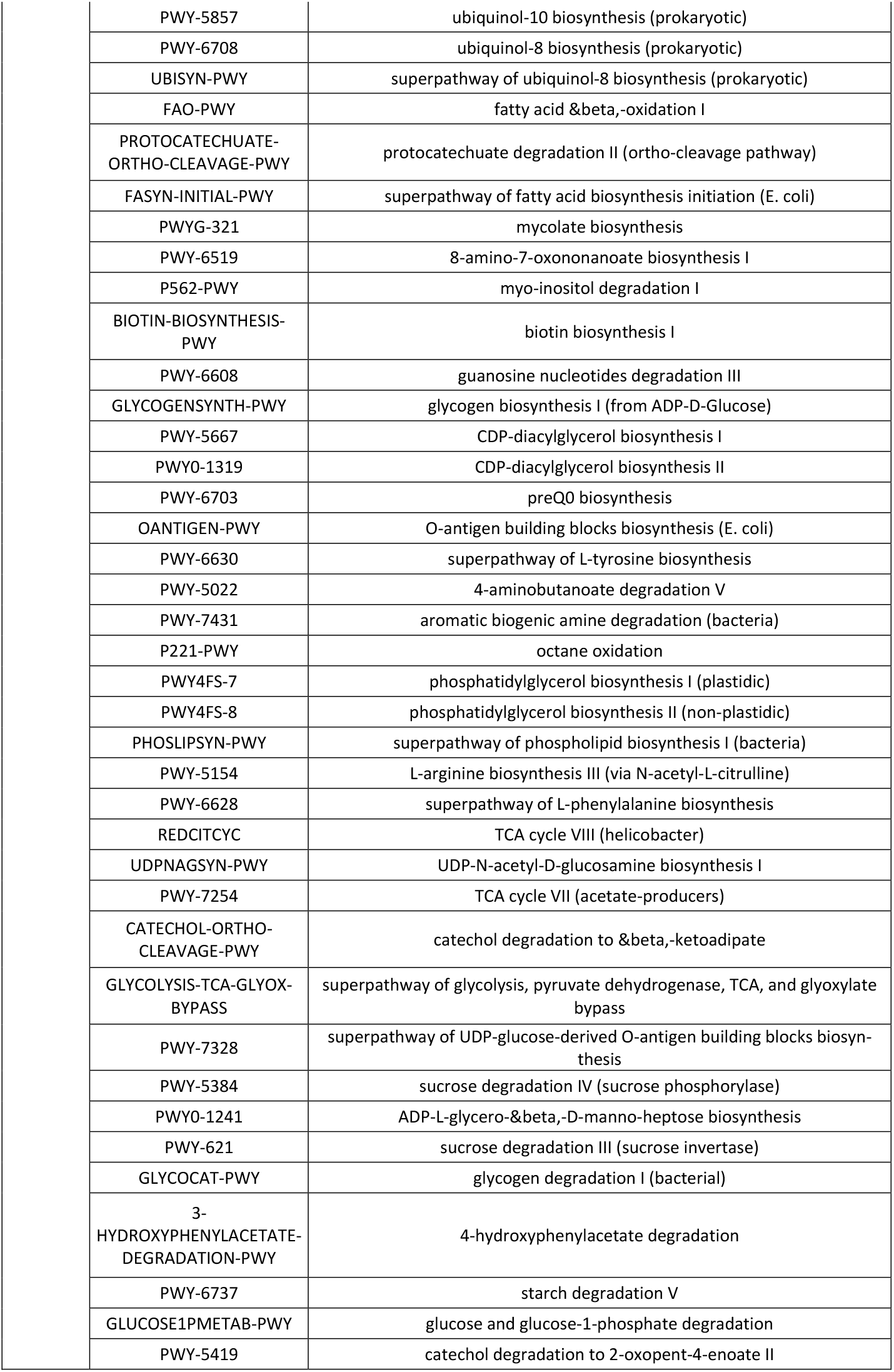
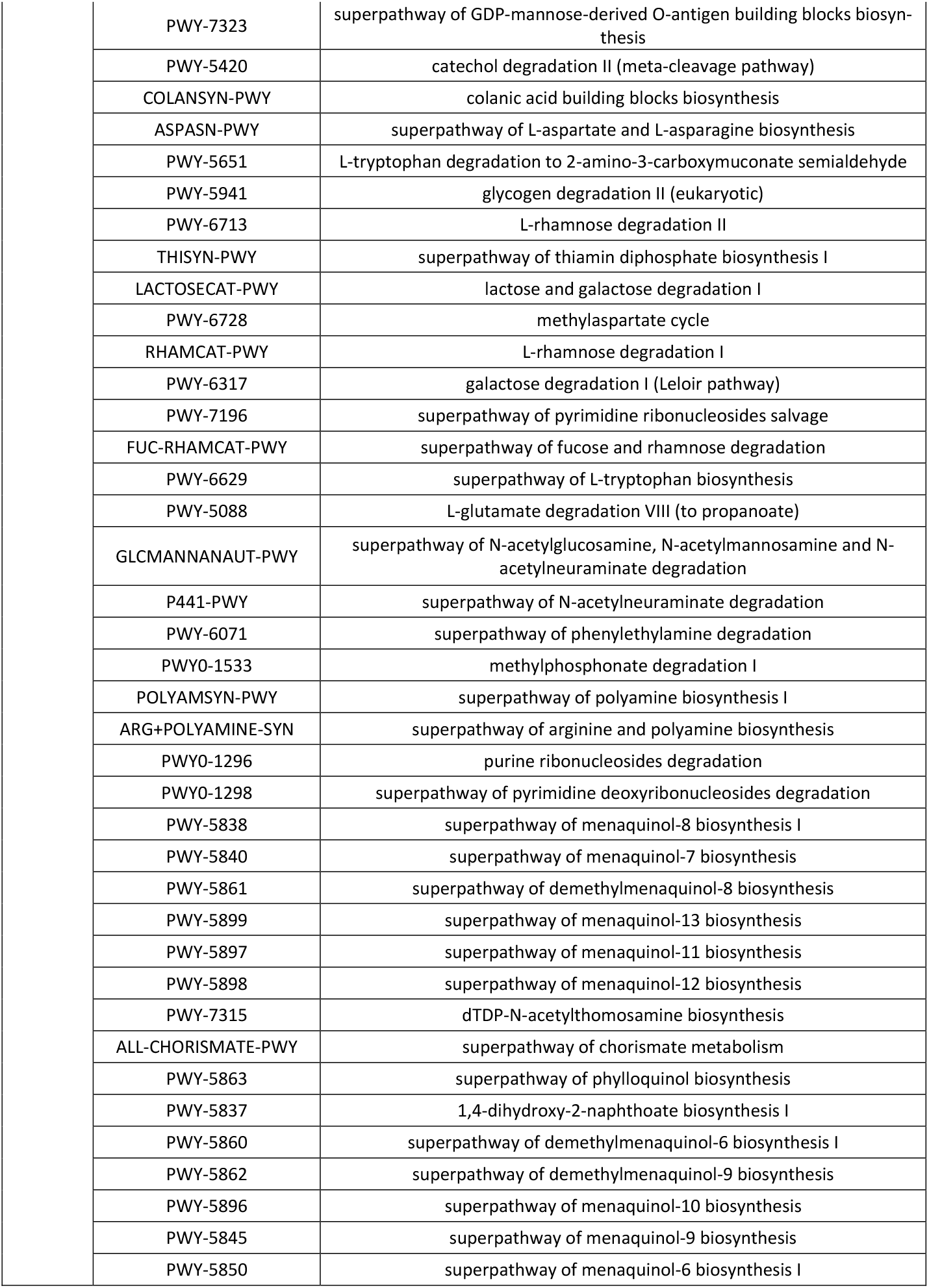
Identification of specific enzymes and pathways, signatures of low score of fibrosis.

On the other hand, six major metabolic pathways were positively associated with the F0 score from both components. One involves the glycolysis and pentose phosphate pathway (PWY-6629), while the 5 others are all involved in the menaquinones and demethylmenaquinones pathway (**Fig 6L**). The low-molecular weight lipophilic components of the cytoplasmic membrane are considered vitamin K_2_ components that is found in most aerobic Gram-positive bacteria and are the main quinones that function as a reversible redox component of the electron transfer chain, mediating electron transfer between hydrogenases and cytochromes. Altogether the functional metagenomics prediction suggests that gram negative bacteria from the Proteobacteria family composed of preQ_0_ biosynthesis and glycolytic pathway are signature of F1-2 fibrosis scores while the vitamin K_2_ biosynthesis pathway from gram negative bacteria such as Actinobateriaceae [31,32] would be signing F0 liver fibrosis score.

## Discussion

We here report a mathematical approach to identify a bacterial 16S rDNA signature in liver tissue and corresponding putative biochemical pathways in patients with low scores of fibrosis. Our main finding is that even low scores of fibrosis (F0 vs F1-2) can be classified by biomarkers from the Proteobacteriaceae family within the liver. The second observation is related to the importance of cohort heterogeneity in term of size and data variability which could be major confounding factors that must be taken into account in multi-centric clinical trials or database. We here present a mathematic approach that could help solving this major and common issue.

A gut metagenomics signature of liver fibrosis in humans has been recently described, suggestive of its causal role in the disease [21]. However, such patients where mostly characterized by a high score of liver fibrosis questioning the putative causal role of the liver microbiota in the disease. We here focused our attention on low scores of liver fibrosis to putatively identify causal factors. We identified mostly gram negative bacteria and notably the Proteobacteria as signature of the F1-2 liver fibrosis scores. Among the families the Proteobacteriaceae, Flavobacteriaceae, and Propionibacteriaceae were discriminating the low fibrosis scores from each other’s. They synthesize LPS, a dramatically inflammatory suggesting a pathophysiological role in development of liver fibrosis, probably via the maintenance of a certain degree of immune vigilance. We further refined our analyses and mostly selected Enterobacteriaceae family from the Proteobacteria phylum suggesting that the liver proinflammation observed during fibrosis would be due or associated with genera from the Enterobacteriaceae family [10]. The Enterobacteriaceae encompass the genera Arthrobacter and Acinetobacter. The mechanisms through which such bacteria could induce inflammation might be linked to the unique structures of their LPS or peptidoglycans [32]. Furthermore, since such bacteria are motile with flagella, one could also contemplate that the flagellar proteins are involved in the liver fibrosis process. However, data report that the TLR5 receptor of flagellin is rather associated with protection against metabolic syndrome, putatively ruling out this hypothesis [33]. Through functional metagenomics production we identified the preQ_0_ biosynthesis pathway as a signature of F1-2 fibrosis scores. Such pathway is notably identified from gram negative bacterial such as Proteobacteriaceae [28,29]. Conversely, the menaquinones and demethylmenaquinones pathways involved in K12 vitamin synthesis were the signature of the F0 score. They are notably produced by the gram positive Actinobacteriaceae such as Bacillus subtilis [31], therefore coherence with our metagenomics findings.

A major hurdle of aggregation of different cohort altogether is related to the heterogeneity of the size of the groups and of the diversity of the variables considered. Regarding invasive analyses such as liver biopsies the group size at completion of the inclusions could be different from what predicted during the calculation of power of the trial. Eventually, the distribution of the variables to be studied could be highly heterogeneous for a given disease. Altogether, we here faced several statistical challenges which are linked to liver fibrosis. The first major step preceding the microbial analysis was a prefiltering and then an adapted pathway to normalize the data to deal with their sparse nature. The package Mixomics [35] used for this study recommends CSS normalization on sparse OTU table counts that could prevent the bias included in the TSS normalization. In addition, it includes multivariate methods for microbiome studies and addresses its limits. In addition, we observed a strong impact of the cohort of origin since the largest cohort from Romania could discriminate the patients from the others based on the 16SrDNA OUT variables. The patients could even be classified by cohort when we used the clinical data as entries showing that this issue also has to be taken into account when analyzing the data. Mathematical approaches to overcome this issue are currently being developed however, little has been done regarding the handling of the 16sRDNA data now widely used by the scientific community that addresses the role of microbiota on diseases and notably liver diseases. Therefore, we here developed several approaches of fairness to overcome the classical cohort impact. Eventually, we noticed that two patients from the F1 groups were distributed with the F2 group. This ectopic distribution could be due to the extreme BMI (>55) featuring a specific clinical phenotype. Conversely, a patient from the F2 group was associated with the F0/F1 distribution. This patient was characterized by his young age (<40 years old) while the mean age of the F2 group was of 54 years old.

The statistical approach required to properly analyze microbial data sets needed to be better fitted to the nature of the data. As a preliminary analysis we performed PCoA since better adapted than PCA to dissimilar and sparse data then followed by a sPLS-DA to identify subsets of 16S rDNA that are discriminatory for the liver fibrosis scores. PLS-DA aims to classify a data set according to the values of a qualitative variable by maximizing the covariance between linear combinations of the observed variables and the qualitative outcome. The sparse version, on the other hand, delivers variables per each component, only selected in the OTU dataset, that are the most discriminatory for the liver fibrosis scores. We focused our attention on the identification of the OTU frequencies within and across each group of patients and on the understanding of the importance that OTUs carry within and across the cohort. We found that the data set is mostly populated by a few high frequency OTUs. However, beside the level of information gained form this approach where overrepresented OTUs we identified cannot rule out that some more information could be obtained from OTUs rarely represented. Therefore, some information could be hidden in the low frequency OTUs. To test this hypothesis we introduced a new normalization approach called TF-IDF [36] originally developed for text mining, to attenuate the effects of the high frequencies OTUs in the data set. Furthermore, aside from the fibrosis scores, it reveals some new predominant taxa at the different taxonomic levels.

In conclusion, the first evidence of the existence of a liver microbiota opens alternate routes for novel therapeutic strategies since specific bacteria could be involved in the process of liver fibrosis. However, to generate information which could serve as a substratum to reach this aim, we here adapted predicted metagenomics and mathematical approaches to the original and novel nature of the tissue metagenomics data set. We here found that these data are constituted of high heterogeneity variables which are dominated by a few high frequency taxa such as Proteobacteria, signature of F1-2 liver fibrosis scores, and Actinobacteria/Firmucutes, signature of F0 liver fibrosis scores. These major taxa are masking information residing in the lower frequency taxa. Predicting metabolic pathways from selected 16S_rDNA-based taxa revealed a role of folate metabolism in F1-2 liver fibrosis scores while a role of vitamin k12 biosynthesis was characterizing F0 liver fibrosis score. Altogether, the combined use of metagenomics, sPLS-DA, TF-IDF and fainess strategies appeared useful since we identified signatures specific to the lower scores of liver fibrosis i.e. at the onset of the disease.

## Acknowledgements

We wish to extend our gratitude to the study participants, investigators, monitors and study nurses who enabled this study. We are grateful to Bogdana Dorcioman MD and the corresponding team from the Laboratory Department and Emergency of the Mures County Hospital, who performed blood analysis and who provided some technical help. This grant was supported by subsides from the Agence Nationale de la Recherche, Novo-Nordisk and Sanofi-Aventis, and the Région Midi Pyrénées to R.B. This research program was partly funded by VAIOMER SAS (project no. 6869/12.06.2014) through the University of Medicine and Pharmacy Tirgu Mures, A subside was allocated to Camille Champion from Institut National des Sciences Appliquées and the Région Occitanie. HT is supported by the excellence initiative VASCage (Centre for Promoting Vascular Health in the Ageing Community), an R&D K-Centre (COMET program - Competence Centers for Excellent Technologies) funded by the Austrian Ministry for Transport, Innovation and Technology, the Austrian Ministry for Digital and Economic Affairs and the federal states Tyrol, Salzburg and Vienna.

## Author contribution

The study was designed by RB, RMN and JML. The experiments and analyses were done by CC, JEC, FG, BL and FS. The Clinical study was designed, surgeries performed by MAR, ME, HT, RMN, DTS, JMF, MF, and JA. The manuscript was written by RB, FB, CC, RMN, JML.

## Conflict of interest

RB and JA receive honorarium from Vaiomer and have shares. BL and FS are employees of Vaiomer.

**Appendix A. Supplementary data**

## Figure and table legends

**Supplementary Figure 1:**
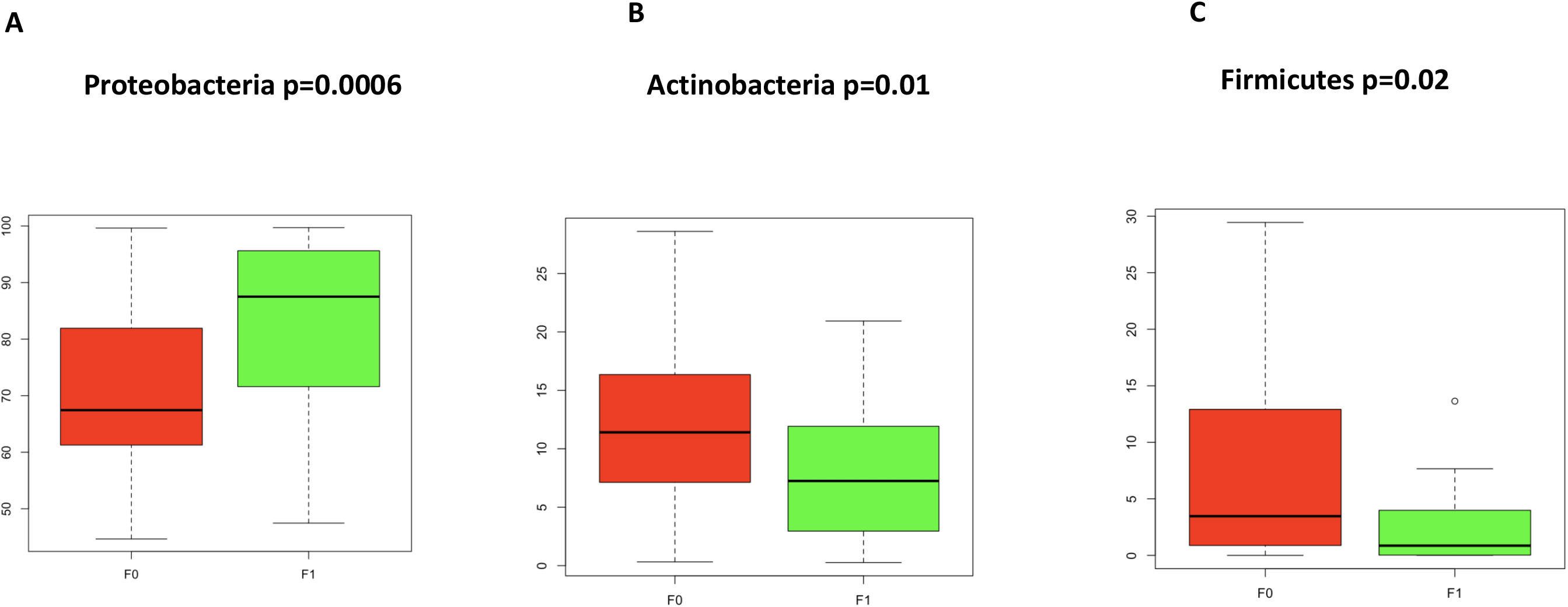
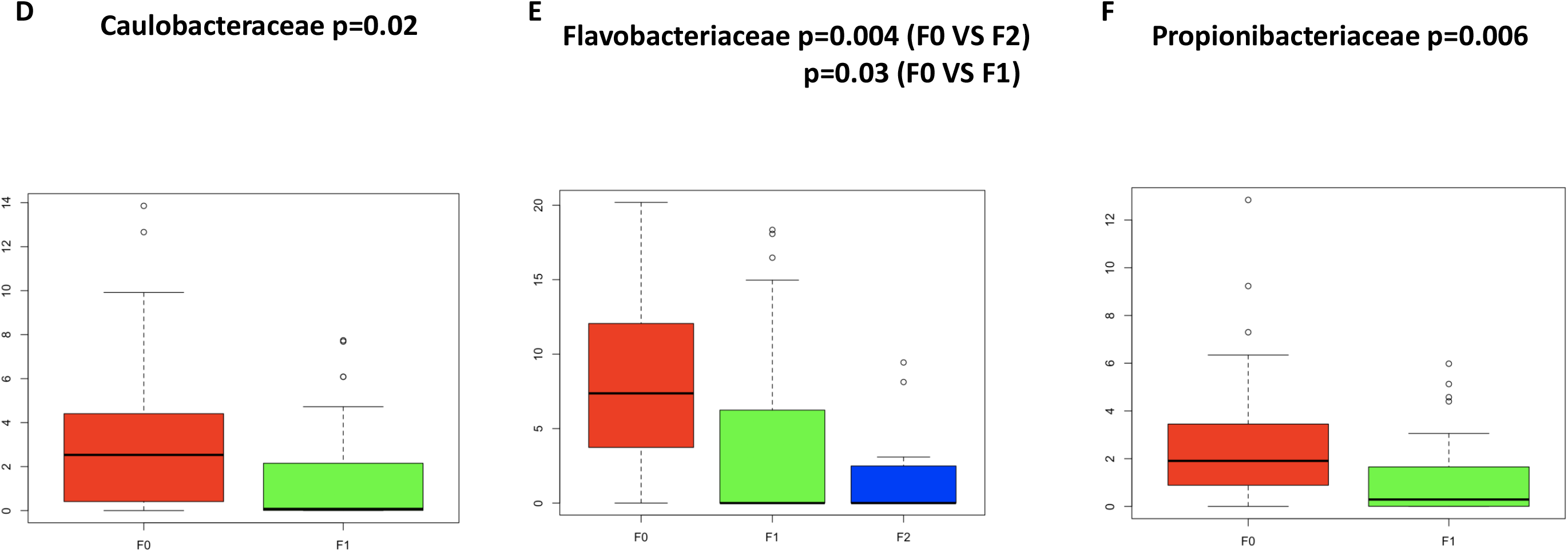
mean frequencies of discriminating taxa. Boxplot representing the frequencies of **A** Proteobacteria, **B** Actinobacteria **C** Firmicutes phyla and **D** Caulobacteraceae, **E**, Flavobacteriaceae, and **F**, Propionibacteriaceae families throughout two groups of liver fibrosis scores (red=F0, green=F1, blue=F2).

**Supplementary Figure 2:**
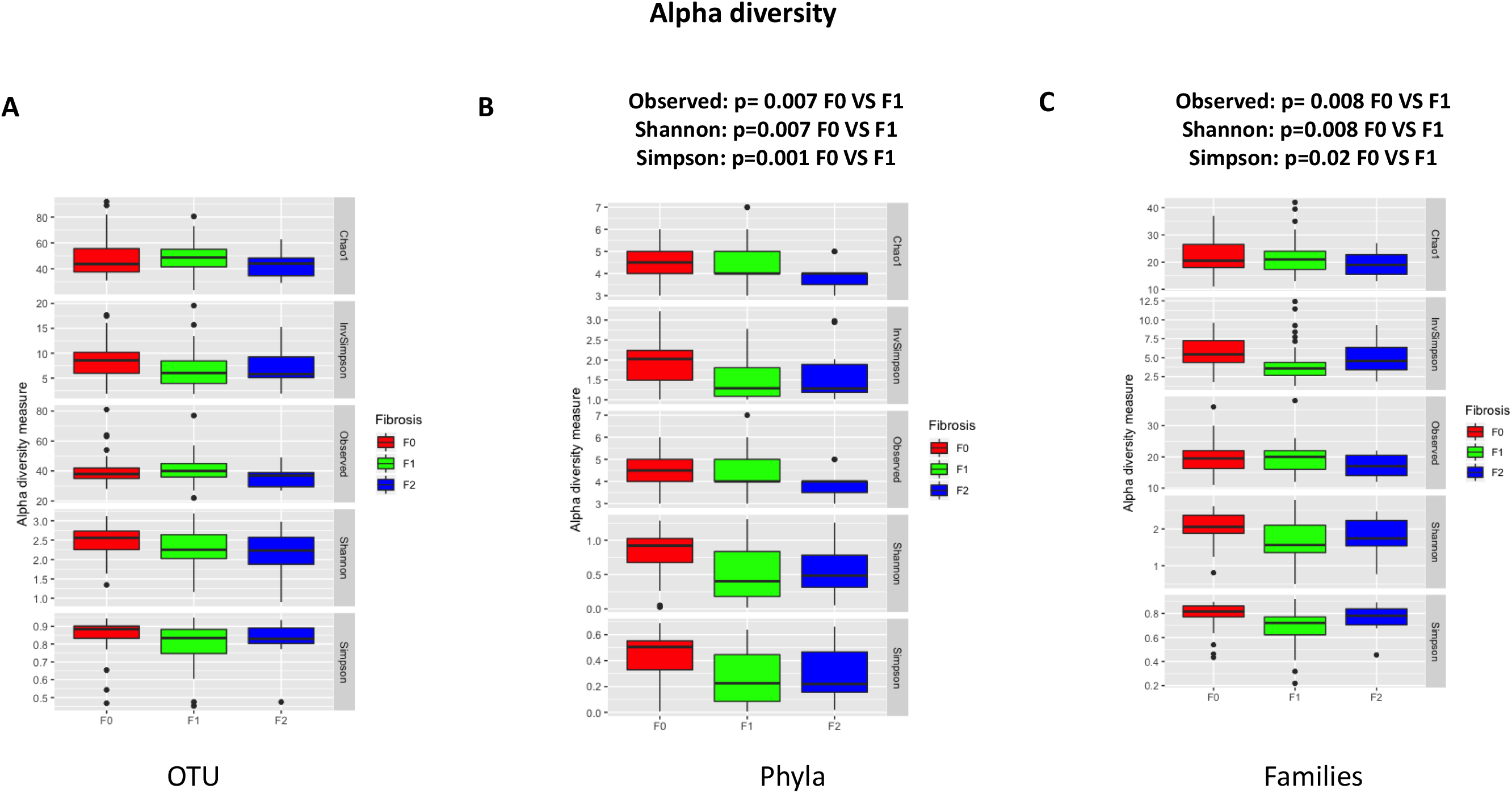
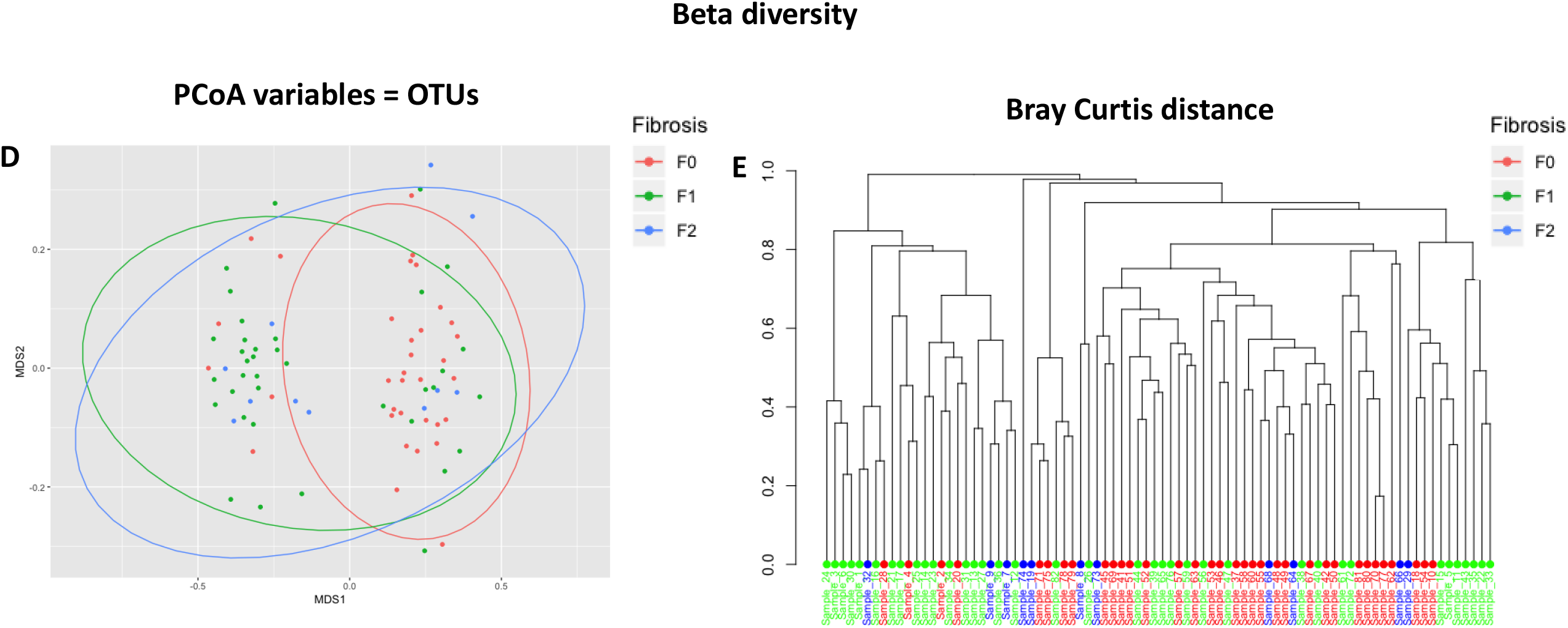
alpha and beta microbial diversity. Boxplot showing microbial alpha diversity **A** at the OTU, **B**, phylum, **C**, and family taxonomic level calculated according to the Chao, Shannon, Simpson, inv Simpson indexes for the 3 liver fibrosis scores. **D** PCoA showing Bray Curtis beta diversity of the normalized OTU table count. Dots are assigned to individual patients and colored according to their fibrosis score (red=F0, blue=F2, green=F1). **E** Hierarchical clustering of patients colored according to their fibrosis score (red=F0, blue=F2, green=F1) based on Bray Curtis OTU distance.

**Supplementary Figure 3:**
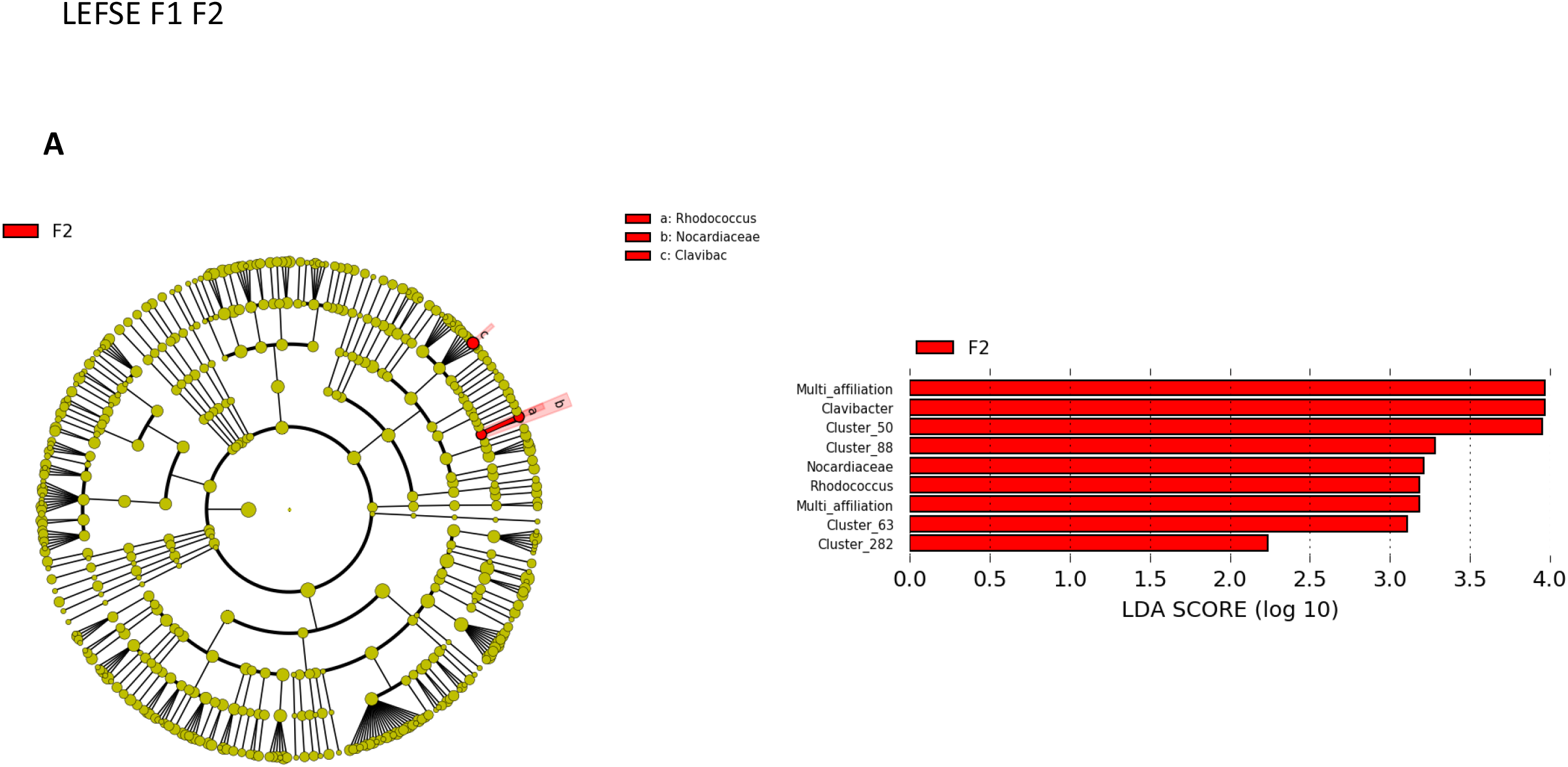
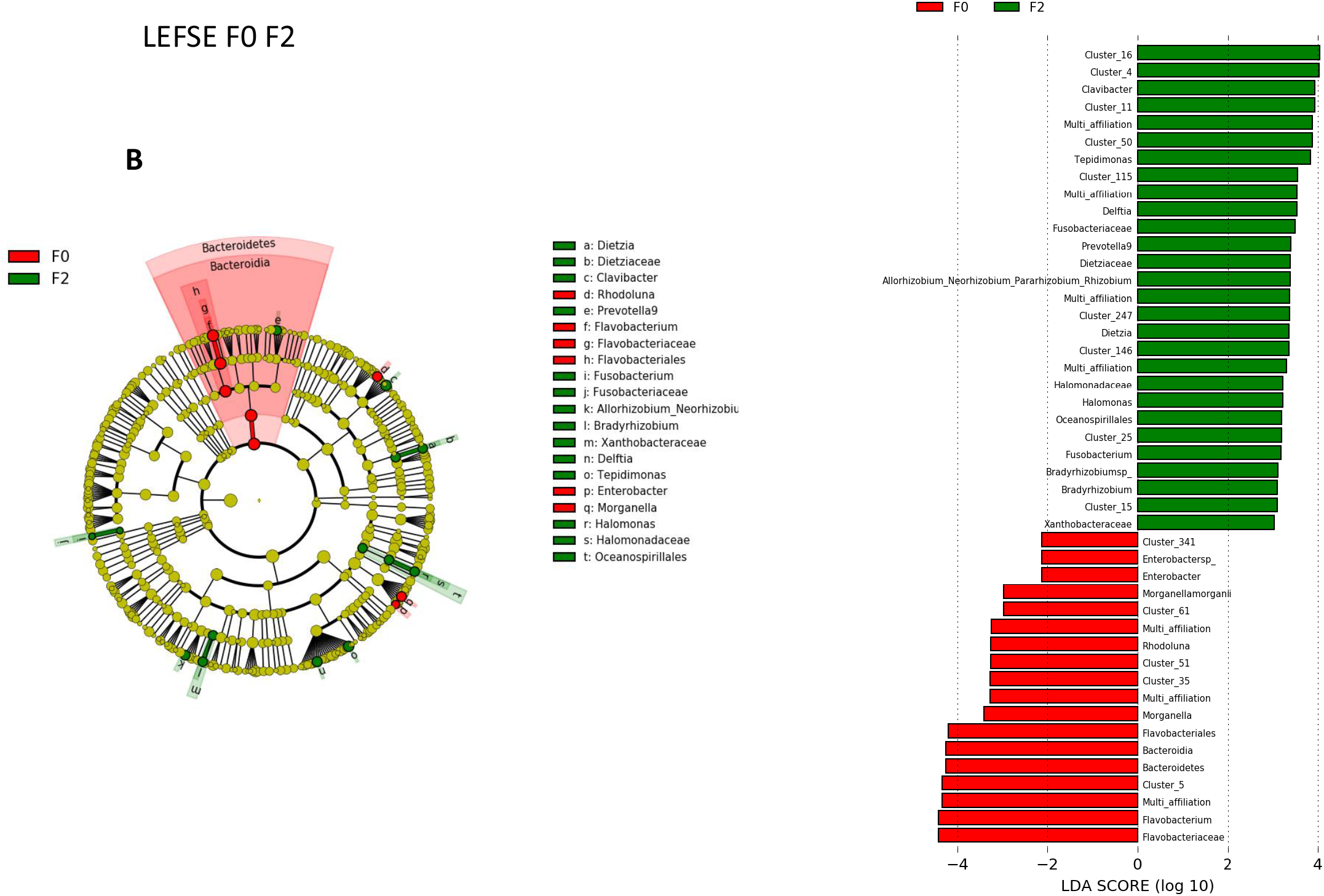
Discriminant microbial signatures identified by linear effect size. LEfSe cladogram and LDA scores of taxonomic assignments from 16S rDNA sequence data of two liver biopsy fibrosis groups **A** F1 vs F2, and **B** F0 vs F2.

